# Aberrant Pain Phenotypes Emerge Following Prenatal Hypoxic-Ischemic Injury in a Rabbit Model of Cerebral Palsy

**DOI:** 10.64898/2026.06.30.735396

**Authors:** Landon T. Genry, Corinne W. Marble, Brendan C. Moline, Patrick J. McGinnis, Cassandra Kramer, Scott Matson, Emily J. Reedich, Elvia Mena Avila, Tracy Santos, Lisa Dowaliby, Natallia Katenka, Marin Manuel, Katharina A. Quinlan, Megan Ryan Detloff

**Author notes:** Corresponding Authors: Megan Ryan Detloff, PhD, Department of Neurobiology & Anatomy, Marion Murray Spinal Cord Research Center, 2900 W. Queen Lane, Philadelphia, PA 19129, Katharina A. Quinlan, PhD, Department of Biomedical and Pharmaceutical Sciences, College of Pharmacy, University of Rhode Island, 130 Flagg Rd, Kingston, RI 02881.

## Abstract

Cerebral Palsy (CP) is the most common motor disability in childhood, and the most frequent comorbidity is pain. Rabbit kits subjected to prenatal hypoxia-ischemia (HI) exhibit allodynia and an expansion of nociceptive afferents in the lumbar spinal cord at postnatal day (P5). In this study, we examined how HI alters the development of multiple sensory modalities and its effect on psychosocial measures and C-fiber distribution in the spinal cord. To do this, we performed an HI surgery to occlude blood flow to fetal New Zealand White rabbits for 40 minutes, or a sham surgery. We performed von Frey, Hargreaves, and a cold allodynia test at P1, P5, P11, and P18. Additionally, we performed open field, a two-texture preference test, and immunofluorescence assays at P18. HI kits exhibit altered development and allodynia in von Frey and Hargreaves and decreased sensitivity to cold. HI kits spend less time on the aversive side of the two-texture preference apparatus and more time in the center of an open field but a higher ratio of that time immobile. This is accompanied by changes in the distribution of C-fibers in the dorsal horn of the cervical and lumbar spinal cord. A principal components analysis revealed prenatal HI produces coordinated alterations across sensory and behavioral domains and multivariate phenotyping captures the overall HI phenotype more comprehensively than any single measure. Overall, HI rabbits kits exhibit altered sensory development, allodynia, anxiety-like behavior, and changes to the distribution of nociceptive afferents in the dorsal horn of the spinal cord.

## Introduction

Cerebral palsy (CP) is an umbrella term for non-progressive movement and postural disorders arising from perinatal injury to the nervous system [17]. CP is the most common physical disability in childhood, with a prevalence of approximately 1-4 per 1000 live births [24; 34], but the lived experience of people with CP extends beyond motor dysfunction [17]. Pain is a major health issue in CP, strongly associated with reduced quality of life [25; 49; 61; 62]. Up to 76% of children and young people with CP experience pain, and about one in three meet criteria for chronic pain [27; 42; 43]. Children with CP report that pain interferes with sleep and their ability to have fun [42] and are over 2.5 times more likely to report psychological symptoms with severe pain [52]. This burden does not diminish with age. A recent meta-analysis of over 1,200 adults with CP reported pain prevalence at 70% [61].

The mechanisms underlying CP-related pain are heterogeneous and incompletely understood. Converging evidence indicates the etiology of chronic pain in CP extends beyond musculoskeletal pathology. Adults with CP exhibit higher prevalence of all pain phenotypes compared to the general population [45]. A subset of individuals with CP report neuropathic qualities such as burning or stabbing sensations, and spinothalamic sensory abnormalities correlate with greater pain intensity, implicating central pain mechanisms [15]. Quantitative sensory testing in children with CP reveals patterns of mechanical and thermal hypoesthesia and mechanical hyperalgesia, suggesting that CNS injury disrupts both discriminative and nociceptive somatosensory processing [7; 48]. This signals the injured nervous system itself is an active contributor to pain experience, not a passive substrate [6; 45].

Despite this, the pathological mechanisms driving altered pain and sensory processing remain poorly characterized. This knowledge gap is consequential: early developmental windows represent critical periods during which aberrant neural connectivity may be amenable to targeted intervention [6; 40]. Progress requires experimental animal models that reproduce the sensory and pain phenotypes in addition to the motor features of CP. The established rabbit model of prenatal hypoxia-ischemia (HI) reliably recapitulates the movement disorder aspects of CP [19; 20; 22; 58], and our group has shown that neonatal HI rabbits exhibit mechanical allodynia and thermal hyperalgesia accompanied by expansion of calcitonin gene related peptide (CGRP)-positive C-fibers in the lumbar spinal cord [46]. Whether this model captures the multimodal sensory dysfunction characteristic of human CP, and whether these changes evolve across the neonatal period in a manner consistent with clinical trajectories has not been systematically examined.

The purpose of this study was to characterize sensory responses across multiple modalities throughout postnatal development in HI rabbits, and to determine whether behavioral changes are associated with reorganization of nociceptive fibers in the dorsal horn at postnatal day (P)18. We hypothesized HI rabbits would exhibit alterations in nociception and psychosocial behavior corresponding to maladaptive changes in afferent distribution within the spinal cord. We found that HI rabbits exhibited hypertonia, altered reflex development, altered nociception, anxiety-like behavior, and altered nociceptive afferent area in the dorsal horn.

## Methods

### Subjects

All experiments were performed in accordance with the United States’ National Institutes of Health’s Guide for Care and Use of Laboratory Animals and approved by the University of Rhode Island’s Institutional Animal Care and Use Committee. Male and female New Zealand White rabbits were used in all experimental groups, and all rabbits were born in-house at the University of Rhode Island. The control group comprised a mix of naïve kits that underwent no surgery (n=12) and kits that underwent sham surgery *in utero* (n=13) which included sedation but no hypoxia-ischemia. Naïve control and sham kits came from 8 litters and were collapsed into a single control group, as separate statistical analyses on each behavioral test revealed inconsequential differences between these conditions, and our primary interest was to examine the effects of the HI surgery compared to uninjured conditions. A comparison of the sham and naïve controls is included in supplemental table 1 and supplemental table 2. The experimental hypoxia-ischemia (HI) group had 24 kits from 3 litters. Two dams provided multiple litters; one dam provided both a sham and an HI litter, and the other provided 1 control litter and 2 sham litters. Kits were monitored at least daily for hydration and visible milk in the stomach. If these factors were unsatisfactory, kits were orally administered supplemental rabbit milk replacer (Wombaroo Food Products, Mount Barker, South Australia). There were 2 stillborn control kits, 2 stillborn sham kits, and 0 stillborn HI kits. Kits were counted as stillborn if they were dead upon birth or died less than 24 hours after birth. Two HI kits died before the 18-day testing period was completed. All rabbits were housed on a 12h light/dark cycle in a temperature and humidity-controlled room in standard Allentown rabbit enclosures. Dams were provided with a nest box lined with hay before their due date. Water, chow (Envigo 2031; Inotiv, Inc., Lafayette, IN), and hay were available *ad libitum* in the home cages.

### Hypoxia-ischemia surgery

Pregnant New Zealand White dams at E22/E23 (∼70% gestation) underwent HI procedures as previously described [20], including sedation with intramuscular administration of ketamine/buprenorphine/dexmedetomidine, (5/0.05/0.025 mg/kg, respectively) or ketamine/xylazine (25/1.5 mg/kg). They were then administered up to 5% isoflurane in O_2_. A Fogarty arterial embolectomy catheter (Edwards Lifesciences Cat # 120602FP or 120403FP) was inserted into the femoral artery up to the level of the bifurcation of the descending aorta. Catheter placement at the bifurcation was confirmed via ultrasonography. The balloon catheter was inflated with saline under continuous Doppler monitoring until blood flow was blocked. This occluded blood flow to the uterine horns and the rabbit fetuses. After 40 minutes, the balloon was deflated and removed. The femoral artery was tied off and the incision was closed. Dams recovered and were returned to their home cages. For sham surgeries, pregnant dams were placed under anesthesia without artery occlusion. Importantly, all dams that underwent surgery recovered and gave birth normally on the expected due date +/- one day.

### Behavioral Testing

All kits underwent repeated behavioral testing at P1, P5, P11, and P18. Care was taken to follow testing order that began with basic weight and length measurements, followed by the open field testing and the righting reflex test. Then, mechanical and hot temperature sensitivities were assessed using von Frey and Hargreaves’, respectively. Hypertonia was determined with the modified Ashworth scale, further modified for rabbits [20]. Afterwards, non-evoked mechanical sensitivity was assessed using the two-texture preference test and cold temperature sensitivity was assessed using a cold allodynia test. von Frey, Hargreaves’, and the cold allodynia test were performed on all paws. The order of paws being tested was randomly determined by the experimenter. For von Frey, Hargreaves’, and cold allodynia tests, kits were allowed to acclimate to each of these testing environments for 5-10 minutes until they were quiet and still but not sleeping. Experimenters remained consistent throughout behavioral testing per litter to limit inter-rater variability. Detailed descriptions of behavioral assessments are outlined below.

### Open Field Test

Behavior in an open field was tested at P18 as rabbit kits reliably have their eyes open by this age [31]. This test was performed in a novel environment, the experimenter would leave the room, and anxiety-related measures were specifically collected [51]. Kits were placed in the center of a large, square open field (1m x 1m x 0.35m) with a soft foam mat lining the floor. Kits freely explored for 5 minutes. The test was recorded with an AUKEY PC-LM3 camera (AUKEY, Shenzhen, China) and analyzed using ANY-Maze software (Stoelting Co., Wood Dale, IL). For video analysis of anxiety-like behavior, the open field was divided into a surround zone which accounts for 68% of the apparatus and a center zone which accounts for 32% of the apparatus using ANY-maze (Stoelting Co., Wood Dale, IL). Signs of anxiety were measured using the amount of time kits spent immobile or in the center of the open field [51; 54]. Immobility detection was set to 65% and the minimum immobility period was set to 2000 ms. We examined measures associated with anxiety-like behavior (time spent, distance traveled, and immobility in the center).

### Righting Reflex

Reflex development was tested using an adapted righting reflex test [20] and was recorded with an AUKEY PC-LM3 camera (AUKEY, Shenzhen, China). Kits were held supine in a box (45cm x 30 cm x 30.5 for P1 and P5 kits, 51 cm x 51 x 30.5 cm for P11 kits, and 1m x 1m x 35 for P18 kits) until experimenters let go of the kits and let the kits right themselves to a prone position. This test was repeated 10 times with at least 10 seconds between each trial. Post-hoc scoring was conducted in a blinded manner to determine righting time and behavioral differences in righting, such as the kit not performing the test in a fluid, continuous, stereotypical movement. Categories included partial righting without complete rotation, disjointed movements, repeated attempts, and attempts that end in falls or loss of balance. These categories constituted struggling. The righting time of all trials for an individual kit at a specific timepoint were averaged, so each kit received an average righting time at each age. Trials were excluded if the video skipped, if the kit was not secured in a supine position before the trial, or if the kit fell asleep during the trial.

### von Frey Test

Mechanical sensitivity was tested using the von Frey test. Testing methodology was adapted from a previous publication from our lab [46]. Rabbits were placed in a chamber with a plastic mesh gridded (8 mm x 8 mm) bottom that allowed access to the plantar surface of the paws of the rabbits. Starting with the 1g filament, a von Frey filament (Stoelting Cat # 58011) was applied perpendicular to the plantar surface of the paw in the ascending method. The filament was applied 5 times to each paw with at least a 10 second pause between each application of the filament [46]. The number of paw withdrawal reflexes was recorded as well as subsequent supraspinal responses, affective responses indicating transmission of sensory information to supraspinal centers. Behaviors that were quantified as supraspinal responses included jumping, rearing, shaking their head/paw/whole body, vocalization, tucking the paw, looking at their paw, kicking, stomping their hindpaws, moving away from the location where the stimulus was applied, an exaggerated response that involved falling over, licking the paw, biting the paw, extension of the limb, holding the paw in the air, scratching their body, or scratching or chewing the plastic grid lining the bottom of their enclosure after application of the filament [14; 64]. The experimenter applied the next weaker filament when responses to 1g are noted, or the experimenter applied the next stronger filament until the kit responded to at least 3 of the 5 applications to one paw or was non-responsive to the 26g filament. In the case the kit was non-responsive to the 26g filament in at least one paw, that paw was given a score of 60g, the force of the next filament in ascending order. The paw withdrawal threshold is considered the lowest von Frey filament that elicits a paw withdrawal response in at least 3 of the 5 applications.

### Hargreaves’ Test

Heat sensitivity was tested using the Hargreaves’ test. Testing methodology was adapted from a previous publication from our lab [46]. Using the Hargreaves apparatus (Model 336G Plantar/Tail Stimulator Analgesia Meter from IITC Life Science, Woodland Hills, CA), the rabbits were placed in a chamber on top of a tempered glass plate. From below, a 4×6 mm noxious radiant light beam was applied to the plantar surface of the paw. The radiant heat increased gradually from rest (20-30 °C) to a maximum of 83 °C at 8 seconds. The paw withdrawal latency was recorded as well as supraspinal responses (listed above). Three trials were collected for each paw and were averaged to yield one score per paw. In addition, the total number of paws tested where the rabbit kit displayed a supraspinal response was quantified [14; 64]. The light beam was automatically shut off at 8 seconds (∼80°C). If the kit did not withdraw its paw within 8 seconds, the paw was labeled as nonresponsive for that trial and was given a score of 9 seconds. A paw was considered “responsive”, if the paw withdrawal latency was < 9 seconds for at least one of its three trials. Hargreaves’ testing was only performed at P1, P5, and P11 because kits’ paws are insulated by fur at P18, and, in our experience, they no longer withdraw from the heat stimulus within 8s.

### Ashworth Test

Categorization of motor deficits was performed using a modified Ashworth scale that was further modified for rabbits as previously described [20]. Briefly, joints in the fore- and hindlimbs were rotated, and their tone was evaluated separately by two experimenters. Kits were considered to be “motor affected” if the muscle tone score was ≥3 in one or more evaluated joint(s) according to both experimenters. After prenatal HI, some kits exhibit hypertonia, while others are unaffected [20].

### Two-Texture Preference Test for Non-Evoked Mechanical Sensitivity

Non-evoked mechanical sensitivity was tested with an adapted two-texture preference test [30] at P18 as they reliably have their eyes open by this age [31]. Rabbits were placed into a square open field apparatus (1m x 1m x 0.35m) with clear plastic walls. Then, the experimenter would leave the room. The floor was half covered by a soft foam mat (same as what was used for open field) and half covered by outdoor floor mats. These outdoor floor mats are a rough, potentially aversive texture with small filaments pointing upwards similar to von Frey filaments or astroturf. The rabbit kit was placed in the center of the open field apparatus and allowed to explore for 5 minutes while being recorded using the same setup as above. From ANY-maze, we extracted measures related to texture preference (time, distance traveled, and immobility on each side).

### “von Freeze” Test for Cold Sensitivity

We designed a new test to assess sensitivity to cold stimuli [37] inspired by the cold plantar assay [10; 11]. In this test, we placed rabbits in a chamber with a plastic mesh gridded (8 mm x 8 mm) bottom that allowed access to the plantar surface of the paws. Once the rabbit was still, a micro-popsicle was gently applied to one of the paws. The application of the micro-popsicle was enough to contact the skin, but not enough to lift or poke the paw. The paw’s withdrawal latency was recorded as well as supraspinal responses (listed above) [14; 64]. In addition, the total number of paws tested where the rabbit kit displayed a supraspinal response was quantified. The paw was labeled nonresponsive if it did not withdraw within 20 seconds and was given a score of 21 seconds.

### Analysis of Nociceptor Distribution via Immunofluorescence

P18 rabbits were euthanized with intraperitoneal injection of Euthasol solution (dosed with 200 mg/kg or more; 390 mg/mL pentobarbital sodium and 50 mg/mL phenytoin sodium; Covetrus, Portland, ME). Transcardial perfusion was performed using chilled 0.01 M phosphate buffered saline (PBS; pH 7.4) until tissue blanching was observed. 4% paraformaldehyde (PFA) in PBS was then infused at a constant flow rate of 10-12 mL/min using a Gilson Minipuls 3 perfusion pump (Gilson Inc., Middleton, WI). The perfused spinal cords were dissected, post-fixed in PFA at 4°C overnight, washed in 0.01M PBS, and then submersed in 30% sucrose in PBS for cryoprotection. C7-C8 and L3-L5 blocks were isolated from the rest of the spinal cord, as they contain sensory afferents that innervate the fore- and hindlimbs, respectively [29; 39; 47]. These blocks were frozen at −80°C until sectioning. For sectioning, spinal cord blocks were embedded in OCT compound (Fisher Scientific, Pittsburgh, PA). Transverse sections were sliced at −20°C using a cryostat (Leica Microsystems, Wetzlar, Germany) and collected in serial at 25 µm thickness. To avoid repeat regions of analysis, every 10^th^ section was used. Sections were mounted on Superfrost plus microscope slides (Thermo Fisher Scientific, Waltham, MA) and stored at −20°C until they were stained.

Sections were labeled with isolectin B_4_ (IB4) and an antibody to detect CGRP to identify non-peptidergic and peptidergic nociceptive afferent fibers, respectively [16]. For staining, sections were dried at 37°C for several hours to improve section adherence to the slide. For antigen retrieval, sections were treated with Dako Antigen Retrieval Solution (Agilent Pathology Solutions, Santa Clara, CA) in a vegetable steamer at 95°C for 10 minutes. Sections were rinsed in 0.01 M oxidized phosphate-buffered saline (OX-PBS; 0.01 M PBS with 0.1 g/L Thimerosal) twice for 5 minutes each and then incubated in a blocking solution (10% normal goat serum, 0.2% Triton X-100 in OX-PBS) for 1 hour at room temperature. This was followed by an incubation in rabbit-to-rabbit blocking reagent (ScyTek Laboratories, West Logan, UT) for 1 hour at room temperature. Sections were then incubated with 1:500 anti-α-calcitonin gene-related peptide rabbit primary antibody (Peninsula Laboratories Cat# T-4032, RRID:AB 518147; BMA Biomedicals, Basel, Switzerland) and Lectin from *Bandeiraea simplicifolia* Biotin Conjugate (Sigma-Aldrich cat. L2140; MilliporeSigma, Burlington, MA) in a solution of 5% normal goat serum and 0.2% Triton X-100 in OX-PBS for 48 hours at room temperature. After three 5-minute washes in OX-PBS, sections were incubated in goat anti-rabbit cross-adsorbed secondary antibody conjugated for Alexa Fluor 488 (Molecular Probes Cat# A-11008, RRID:AB_143165; Thermo Fisher Scientific, Waltham, MA) and streptavidin conjugated for Alexa Fluor 594 (Jackson ImmunoResearch cat. 016-580-084; Jackson ImmunoResearch Labs, West Grove, PA) for 2 hours at room temperature. After three more 5-minute washes in OX-PBS, sections were cover slipped (Corning Cat No. 2990-245, Corning, NY) using DAPI Flouromount-G (Southern Biotech, Birmingham, AL) mounting medium. After cover slipping, slides were stored at room temperature in microscope slide boxes.

Stained sections were imaged at 20x (Leica DM6 B microscope; Leica Microsystems, Wetzlar, Germany). Depending on tissue availability, two to three representative images of the right and left dorsal horn of each animal were imaged. The dorsal root was visible in all images to ensure consistent quantification at the entrance of the roots. The proportional area of CGRP and IB4 were calculated within the cap of the dorsal horn (laminae I-II) using a moment-preserving thresholding algorithm [60] available through the Auto Threshold plugin on ImageJ (NIH, Bethesda, MD). The proportional area of colocalized CGRP- and IB4-positive area was determined using the Colocalization plugin on ImageJ (NIH, Bethesda, MD). The proportional area of CGRP in the deep dorsal horn (lamina III) was calculated with the same method. Representative images for publication were imaged at 40x with a Stellaris 5 confocal (Leica Microsystems, Wetzlar, Germany). Samples were excluded from analysis if the tissue could not be recovered for staining. 3 HI kits were excluded from cervical analysis, and 3 HI, 3 control, and 1 sham kit were excluded from lumbar analysis. Values from the analyzed sections of each dorsal horn were averaged and the average of the right dorsal horns and the average of the left dorsal horns are represented individually.

### Data Analysis and Statistics

All graphical analyses and statistical analyses for righting reflex, two texture preference, open field, and immunofluorescence were performed using GraphPad Prism 10.6.1 (Dotmatics, Boston, MA). Data analyzed using GraphPad Prism at multiple timepoints (righting reflex) were analyzed using a mixed effects model with a Geisser-Greenhouse correction. Tukey’s multiple comparisons test was used to analyze the difference between control and HI kits at each timepoint. Data analyzed at singular timepoints (two-texture preference test, open field test anxiety measures, immunofluorescence, and principal components) was assessed for normality using QQ plots and the Shapiro-Wilk test. Data that passed normality was analyzed with a Welch’s t-test. Data that did not pass normality was analyzed with a non-parametric Mann-Whitney test.

Disparities in behavioral measurements from each limb or side of the spinal cord occur in people with CP [13] as well as in our model [20]. Thus, data acquired from each paw or dorsal horn was treated as an individual data point rather than an average of the two sides together to generate a single value for each kit. von Frey, Hargreaves’, and von Freeze were analyzed with the RStudio (version 2023.06.0+421) packages lme4, lmerTest, and emmeans to allow for clearer interpretation of changes over time in our longitudinal models. We performed linear mixed effects analyses with fixed effects for age and group and a random intercept for individual paws of rabbits to account for the repeated measurements on each paw from P1 to P18. Age was centered, so group effects were estimated at P1. In some cases, visual analysis of graphed means warranted testing linear spline mixed effects models with knots at specific time points. In these cases, initial models to test model fit were performed using maximum likelihood estimation. Models were compared by running an ANOVA and examining AIC values. If models were significantly different, the model with the lowest AIC value was chosen. When there was no statistically significant difference between the models, the simpler linear mixed effects model without the knot term to account for the change in slope was chosen. Linear spline mixed effects models were used in the von Frey forepaws model with a knot (the inflection point) at P11 and the Hargreaves hindpaws model with a knot at P5. In these cases, an additional age term was added to account for the change in slope after the knot. After choosing the best model, analyses were performed using restricted maximum likelihood estimation. From these models, we could extract the change in response for each one unit increase in age, the effect of HI on responses at P1, and the effect of HI on the change in response over time. To assess group differences, estimated marginal means were calculated and pairwise comparisons between groups at each timepoint were performed with Bonferroni’s adjustment to the p-values [26].

To explore the relationships amongst the immunofluorescence and behavioral data, principal component analysis (PCA) was performed using the R (version 4.5.2) package syndRomics [59].The dataset used for PCA included 1) von Frey, Hargreaves’ and von Freeze pain scores at P18 with all paws averaged per animal, 2) Deep and superficial dorsal horn CGRP+ fiber area, IB4+ fiber area, and colocalization area of CGRP+ and IB4+ fibers, with all paws averaged per animal, 3) the ratio of time spent and immobility between the center and surround in the open field test and 4) the ratio of time spent and immobility on the aversive side versus the non-aversive side of the two-texture preference assay. Data was imputed and then standardized so that the mean was 0 and the standard deviation was 1 to ensure that each variable contributed equally to the analysis. Group differences in PC scores were analyzed using Welch’s t test.

### Data Sharing

Study data are deposited at the Harvard Dataverse and available for download at https://doi.org/10.7910/DVN/4MYRSF. Experimental protocols and detailed methods can be found with the dataset, and cold allodynia behavior protocol can be found on protocols.io (https://dx.doi.org/10.17504/protocols.io.eq2lym31wlx9/v1). Any additional information will be provided to researchers upon request.

## Results

### HI kits exhibit motor deficits after prenatal HI

Motor deficits were evaluated in rabbits at each timepoint. Hypertonia was assessed with Ashworth scoring tailored to rabbits [20]. Of the rabbits exposed to prenatal HI, 54% were considered motor affected (**Figure 1A**), consistent with previous studies [20; 53].

**Figure 1:**
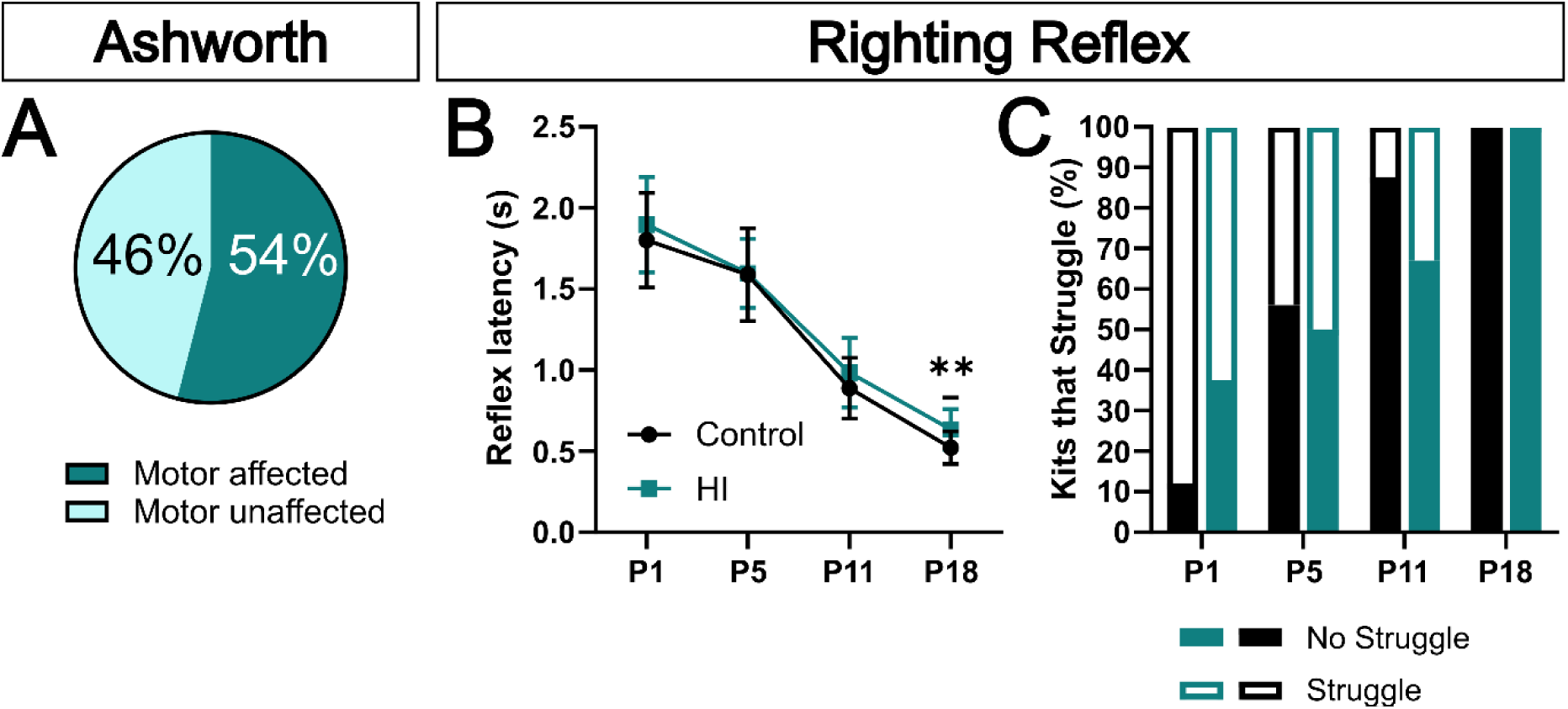
HI rabbit kits exhibit hypertonia but no changes to righting reflex compared to controls. (**A**) Percentage of kits that were motor affected. (**B**) Average righting reflex latency over time. Control and HI kits exhibited similar righting times at P1 (p=0.2583), P5 (p=0.9092), and P11 (p=0.1075). However, at P18 HI kits exhibited slower righting reflex latencies (0=0.0011). (**C**) The percentage of trials that were considered successful (without struggle) at each time point in control and HI kits. *Statistics:* Control N = 25, HI N = 24, (B) Mixed-effects analysis (Group: p=0.0559, F=3.843, Age: p<0.0001, F=390.8, Group X Age: p=0.5634, F=0.6342). Data in (B) represented as mean +/- SD. **p<0.01

Kits were timed in their ability to right themselves to prone from a supine position. Righting reflex latency significantly decreased with rabbit age (Age: F=390.8, p<0.0001). There was no main effect of Group (F=3.853, p=0.0559), and there was not a significant interaction (Group X Age: F=0.5643, p=0.6342) (Figure 1B). Post-hoc comparisons showed control and HI kits took the same amount of time to right themselves at P1 (p=0.2583), P5 (p=0.9092), P11 (p=0.1075), but P18 HI kits took longer (p=0.0011) (Figure 1B). A normal righting reflex is observed to be a single, smooth, continuous rolling motion to right their body from supine to prone position, beginning with their head. Kits that did not display the smooth continuous motion were noted to “struggle.” At P1, the majority of kits struggled to right themselves, including 62.5% of HI kits and 88% of control kits (Figure 1C). With increasing age, fewer kits struggled to right themselves until, at P18, all of the control and HI kits performed the righting reflex in a single, continuous, smooth rolling motion beginning with their head. These results indicate a similar righting reflex development over time in control and HI kits with HI kits’ reflex being slightly slower at P18.

### HI kits display behavior consistent with altered mechanical nociceptive development

After prenatal HI, the developmental trajectory of mechanical nociception is altered in HI kits. In control kits, there was an overall increase in mechanical sensitivity from P1 to P18: on average, there was a 2.31g decrease in forepaw withdrawal thresholds each postnatal day from P1 to P11 (Age: β=-2.31, p<0.0001). After P11, there was a change in their developmental trajectory, and paw withdrawal thresholds only decreased by an average of 0.28g each day until P18 (Age after P11: β=2.03, p=0.0042). In contrast, neonatal HI kits had an average paw withdrawal threshold 20.02g lower than controls (Group: β=-20.02, p<0.0001) at P1. As HI kits aged, from P1 to P11, there was only a slight increase of 0.21g/day in average paw withdrawal thresholds (Group X Age: β=2.52, p<0.0001). After P11, HI kits had an average decrease in paw withdrawal thresholds each day of 1.4g (Group X Age after P11: β=-3.64, p=0.0003) until P18, which is steeper than controls (**Figure 2A**). Pairwise comparisons of estimated marginal means revealed HI kits had lower average paw withdrawal thresholds than control kits in the forepaws at P1 (P<0.0001) and P5 (p<0.0001) but not P11 (p=0.4192) and P18 (p>0.9999) (Figure 2A). This result seems to be partially driven by the differences in the percentage of kits that were nonresponsive to the von Frey filaments in control and HI kits’ forepaws, particularly at P1 (Figure 2B). The lack of significant group differences at P11 and 18 ages could be impacted by a floor effect brought about by the increasing mechanical sensitivity with age. The percentage of forepaws where application of a von Frey filament elicited supraspinal responses was similar in both groups (Figure 2C).

**Figure 2:**
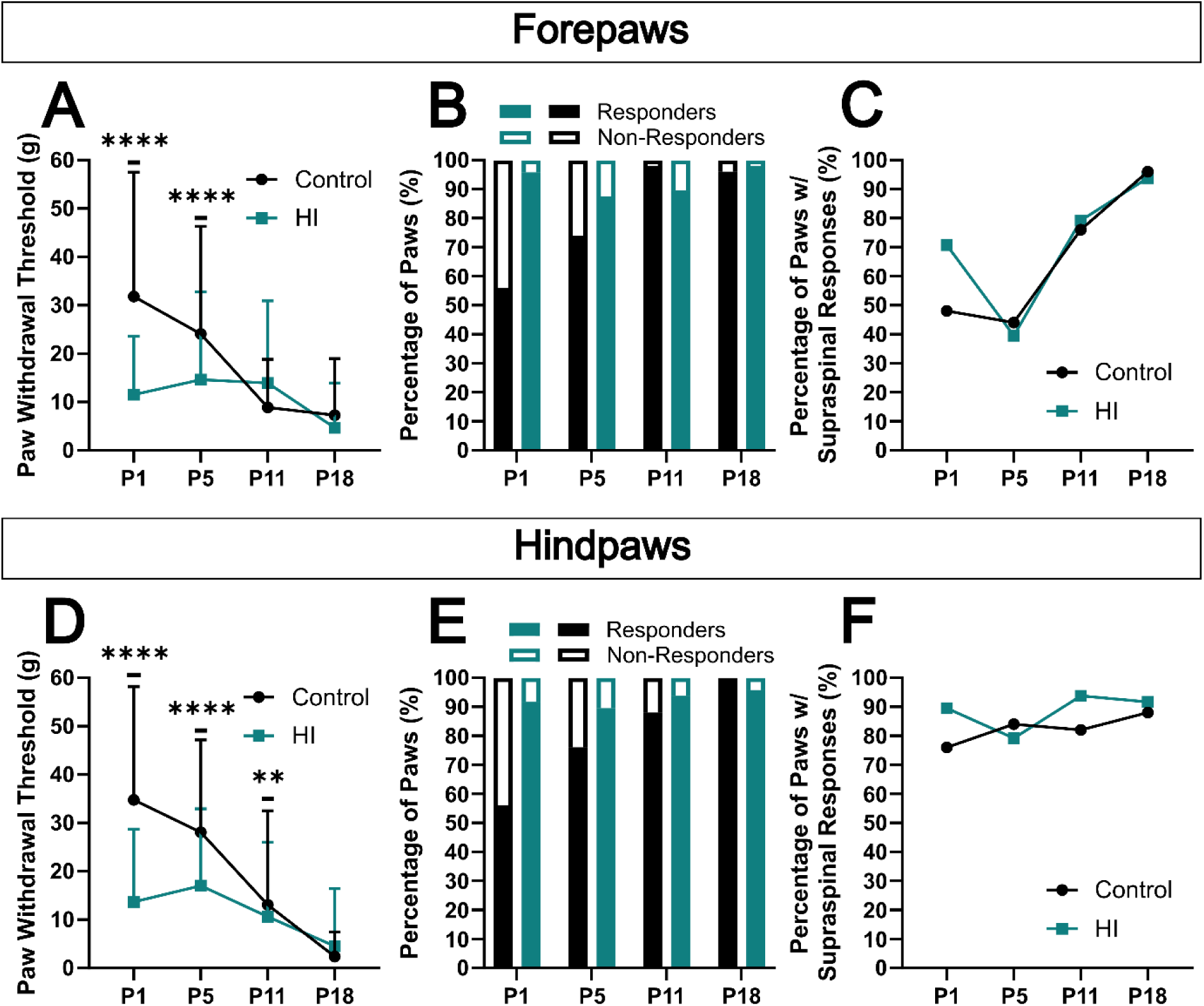
HI rabbit kits are hypersensitive to mechanical stimuli at P1, P5, and P11 in the von Frey test. (**A**) Average forepaw withdrawal thresholds plotted across time. HI kits exhibited lower forepaw withdrawal thresholds compared to control kits at P1 (p<0.0001) and P5 (p<0.0001), but not P11 (p=0.4192) or P18 (p>0.9999). (**B**) Percentage of forepaws that were considered responders (withdrew at least 3/5 times at 26g or before) or nonresponders (did not withdraw at least 3/5 times during testing) at each time point in control and HI kits. (**C**) The percentage of forepaws where the kit exhibited at least one supraspinal response during testing plotted across time. (**D**) Average hindpaw withdrawal thresholds plotted across time. HI kits exhibited lower hindpaw withdrawal thresholds compared to control kits at P1 (p<0.0001), P5 (p<0.0001), and P11 (p=0.0.0046) but not P18 (p=0.1516). (**E**) Percentage of hindpaws that were considered responders or nonresponders at each time point in control and HI kits. (**F**) The percentage of hindpaws where the kit exhibited at least one supraspinal response plotted across time. *Statistics:* Control N = 50 paws, HI N = 48 paws, (A) Linear spline mixed effects model (Group: β=-20.02, p<0.0001, Age: β =-2.31, p<0.0001, Age after P11: β=2.03, p=0.0042, Group X Age: β =2.52, p<0.0001, Group X Age after P11: β=-3.64, p=0.0003). (D) Linear mixed effects model (Group: β=-18.42, p<0.0001, Age: β=-1.96, p<0.0001, Group X Age: β=1.33, p<0.0001). Data in (A) and (D) represented as mean +/- SD. **p<0.01, ****p<0.0001.

In the hindpaws, the results are similar. On average, from P1 to P18, control kits had a 1.96g/day decrease in their paw withdrawal thresholds (Age: β=-1.96, p<0.0001). HI kits were initially more sensitive in their hindpaws with an average paw withdrawal threshold at P1 that was 18.42g lower than controls (Group: β=-18.42, p<0.0001); however, the rate of decline in paw withdrawal thresholds in HI kits was slower, only a 0.63g/day (Group X Age: β=1.33, p<0.0001) average decrease (Figure 2D). Post-hoc comparisons revealed HI kits had lower average hindpaw withdrawal thresholds at P1 (p<0.0001), P5 (p<0.0001), and P11 (p=0.0046) but not P18 (p=0.1516) (Figure 2D). Again, the difference in paw responsiveness seems to contribute to these differences, particularly at younger ages (Figure 2E). In contrast to the forepaws, the percentage of hindpaws where application of the von Frey filament produced a supraspinal response in the kit was similar between control and HI kits over time (Figure 2F).

Together, these results indicate that typical rabbit nociceptive development involves an increase in mechanical sensitivity accompanied by variability in the awareness of stimulation and the affective response over time. The changes to mechanical nociceptive development after prenatal HI point towards accelerated development leading to mechanical allodynia at several timepoints.

### Prenatal HI is associated with thermal hyperalgesia and altered thermal nociceptive development

HI kits exhibited altered development of thermal nociception in response to heat. In control kits, the average forepaw withdrawal latency did not change from P1 to P11 (Age: β=-0.04, p=0.1303). However, this was not true for HI kits. Although there was no difference in the control and HI kits’ average forepaw withdrawal latencies at P1 (Group: β=-0.006, p=0.9805), HI kits did have an average decrease in their forepaw withdrawal latencies of 0.14s/day (Group X Age: β=-0.10, p=0.0128) from P1 to P11 (**Figure 3A**). This difference in the developmental trajectory of control and HI kits’ forepaw withdrawal latencies is confirmed by post-hoc pairwise comparisons. There was no difference between control and HI kits’ forepaw withdrawal latencies at P1 (p=0.9805) or P5 (p=0.0544); however, HI kits had a shorter forepaw withdrawal latency at P11 (p=0.0008) (Figure 3A). This change in sensitivity is not related to the percent of non-responsive control and HI kits’ forepaws, since we can observe that the great majority of all kits were responsive to heat (Figure 3B). There were no differences in the percentage of forepaws where the application of the heat stimulus caused the rabbit kits to display supraspinal responses (Figure 3C).

**Figure 3:**
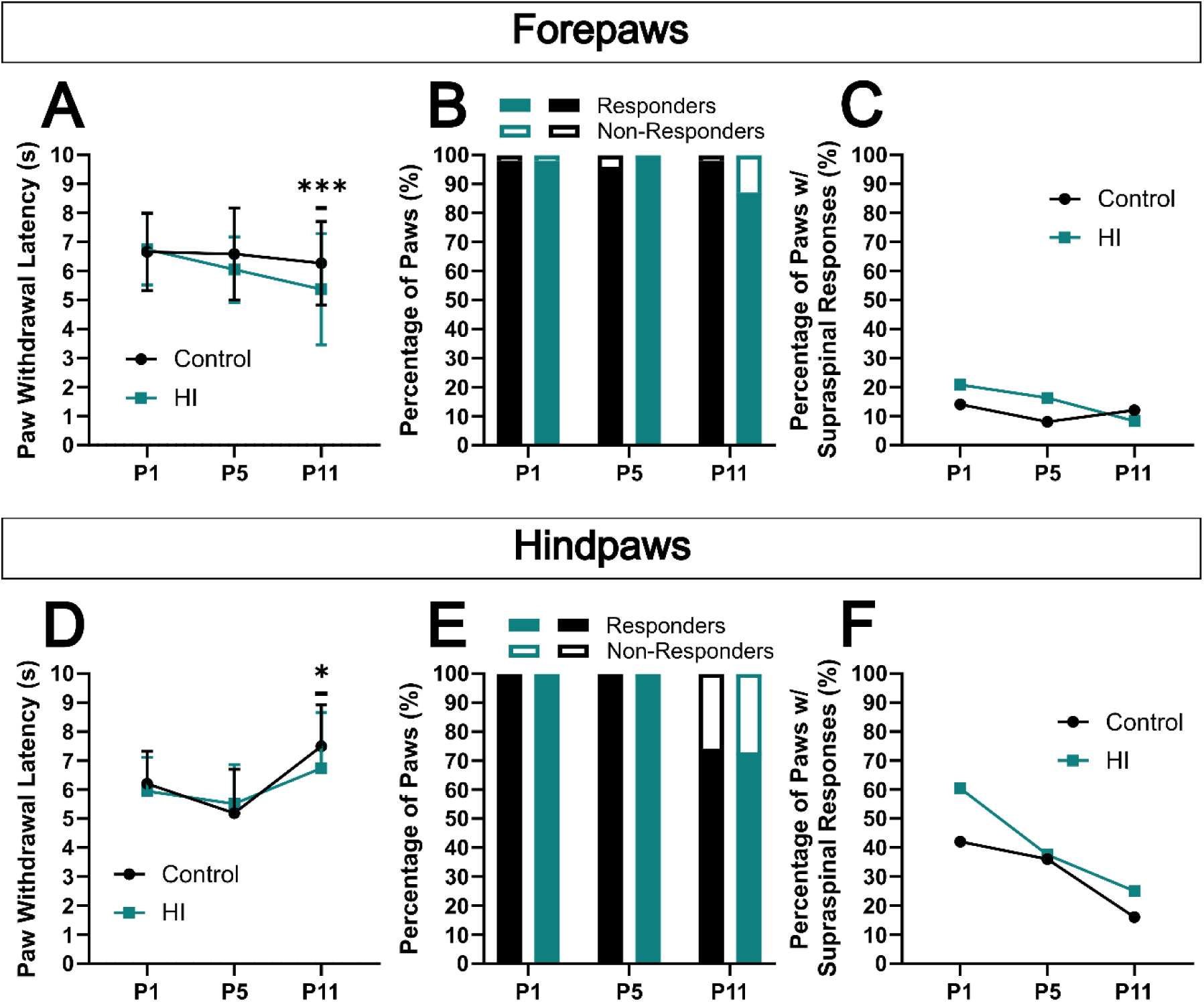
HI rabbit kits are hypersensitive to heat stimuli in the Hargreaves’ test at P11. (**A**) Average forepaw withdrawal latencies plotted across time. HI kits exhibited a lower forepaw withdrawal latency compared to control kits at P11 (p=0.0008) but not P1 (p=0.9805) or P5 (p=0.0544). (**B**) Percentage of forepaws that were considered responders (withdrew at 8s or before) or nonresponders (did not withdraw at 8s or before) at each time point in control and HI kits. (**C**) The percentage of forepaws where the kit exhibited at least one supraspinal response during testing plotted across time. (**D**) Average hindpaw withdrawal latencies plotted across time. HI kits exhibited a lower hindpaw withdrawal latency compared to control kits at P11 (p=0.0281), but not P1 (p>0.9999) or P5 (p=0.7604). (**E**) Percentage of hindpaws that were considered responders or nonresponders at each time point in control and HI kits. (**F**) The percentage of hindpaws where the kit exhibited at least one supraspinal response plotted across time. *Statistics:* Control N = 50 paws, HI N = 48 paws, (A) Linear mixed effects model (Group: β=-0.006, p=0.9805, Age: β=-0.04, p=0.1303, Group X Age: β=-0.10, p=0.0128). (D) Linear spline mixed effects model (Group: β= −0.26, p=0.3633, Age: β=-0.26, p=0.0003, Age after P5: β=0.64, p<0.0001, Group X Age: β=0.15, p=0.1330, Group X Age after p5: β=-0.33, p<0.0222). Data in (A) and (D) represented as mean +/- SD. *p<0.05, ***p<0.001

Consistent with the forepaws, there were no differences between control and HI hindpaw withdrawal latencies at P1 (p>0.9999) and P5 (p=0.7604), but HI kits had a shorter average hindpaw withdrawal latency at P11 (p=0.0281) (Figure 3D). In the hindpaws of control kits, the average withdrawal latency decreased 0.26s/day between P1 and P5 (Age: β=-0.26, p=0.0003), and from P5 to P11, it increased an average of 0.38s/day (Age after P5: β=0.64, p<0.0001). There was no difference in the average age-related decrease from P1 to P5 (Group X Age: β=0.15, p=0.1330) in control and HI kits. However, from P5 to P11, HI kits had a smaller age-related increase compared to controls; HI kits’ withdrawal latencies increased an average of 0.20s/day from P5 to P11 (Group X Age after P5: β=-0.33, p=0.0222) (Figure 3D). The percent of nonresponsive hindpaws and supraspinal responses decreased with postnatal age (Figure 3E-F) and do not follow the same pattern as hindpaw withdrawal latencies (Figure 3A). At P1 and P5, 100% of both control and HI hindpaws were responsive (Figure 3E), but this decreased to 74% of control hindpaws and 73% of HI hindpaws at P11. Supraspinal responses were evoked when stimulating 42% of control and 60% of HI hindpaws at P1 (Figure 3F). This decreased at P5 to 36% of control hindpaws and 38% of HI hindpaws and even further to 16% of control hindpaws and 25% of HI hindpaws at P11 (Figure 3F).

Taken together, these data illustrate prenatal HI is associated with an altered developmental trajectory of thermal nociception leading to the development of heat-evoked thermal hyperalgesia by P11.

### Prenatal HI has limited effects on cold sensitivity

When examining the developmental curves of cold sensitivity using von Freeze, there were no differences between control and HI. In the forepaws, control kits had a 0.29s average increase/day in withdrawal latency (Age: β=0.29, p<0.0001) from P1 to P18. There were no differences in the average age-related change in withdrawal latency in the forepaws between HI and control kits (Group X Age: β= −0.07, p=0.490) (**Figure 4A**). After pairwise post-hoc comparisons, there were no differences in forepaw withdrawal latencies between control and HI kits at P1 (p=0.4952), P5 (p=0.2260), P11 (p=0.0683), and P18 (p=0.1085) (Figure 4A). Similar percentages of forepaws were considered to be responsive between control and HI rabbit kits (Figure 4B). The percentage of paws where stimulation with the micro-popsicle elicited supraspinal responses in the kit was relatively low in both control and HI kits (Figure 4C).

**Figure 4:**
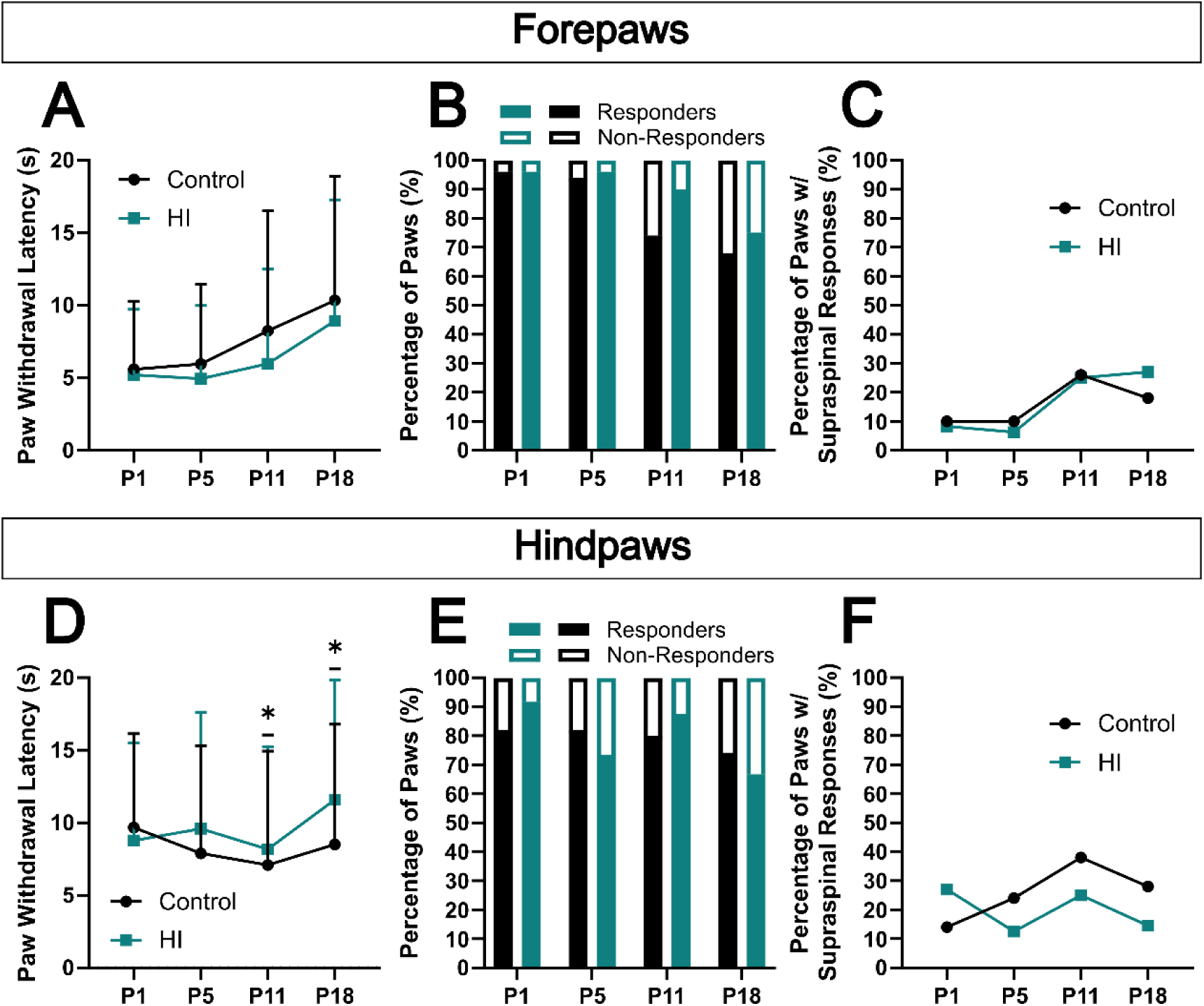
There are minor differences between controls and HI in response to cold stimuli in the von Freeze test. (**A**) Average forepaw withdrawal latencies plotted across time. HI kits exhibited similar forepaw withdrawal latencies compared to control kits at P1 (p=0.4952), P5 (p=0.2260), and P11 (p=0.0683), and P18 (p=0.1085). (**B**) Percentage of forepaws that were considered responders (withdrew at 20s or before) or nonresponders (did not withdraw at 20s or before) at each time point in control and HI kits. (**C**) The percentage of forepaws where the kit exhibited at least one supraspinal response during testing plotted across time. (**D**) Average hindpaw withdrawal latencies plotted across time. HI kits exhibited longer hindpaw withdrawal latencies compared to control kits at P11 (p=0.0473) and P18 (p=0.0261), but not P1 (p=0.8482) and P5 (p=0.5591). (**E**) Percentage of hindpaws that were considered responders or nonresponders at each time point in control and HI kits. (**F**) The percentage of hindpaws where the kit exhibited at least one supraspinal response plotted across time. *Statistics:* Control N = 50 paws, HI N = 48 paws, (A) Linear mixed effects model (Group: β=-0.73, p=0.495, Age: β=0.29, p<0.0001, Group X Age: β=0.-0.07, p=0.490). (D) Linear mixed effects model (Group: β=-0.23, p=0.848, Age: β=-0.06, p=0.470, Group X Age: β=0.19, p=0.107). Data in (A) and (D) represented as mean +/- SD. *p<0.05

In the hindpaws, there were no changes to withdrawal latencies over time in the controls (Age: β=-0.06, p=0.470) and no difference in the age-related change of HI kits compared to controls (Group X Age: β=0.19, p=0.107) (Figure 4D). Post-hoc pairwise comparisons revealed there were no differences between control and HI hindpaw withdrawal latencies at P1 (p=0.8482) and P5 (p=0.5591), but HI kits had higher hindpaw withdrawal latencies at P11 (p=0.04773) and P18 (p=0.0261) (Figure 4D). Control and HI kits had a similar percentage of hindpaws considered to be responsive (Figure 4E). The percentage of hindpaws where stimulation produced supraspinal responses from the kits was small in both control and HI kits throughout development (Figure 4F). These data indicate prenatal HI may lead to less cold sensitivity over time.

### Prenatal HI is associated with increased non-evoked mechanical sensitivity at P18

In addition to testing evoked mechanical sensitivity using von Frey hairs, we also tested non evoked mechanical sensitivity using a two-texture preference test, where one side of an open field was covered with a rough, outdoor floor mat (which we predicted to be aversive), while the other side was covered with a soft foam mat (which we predicted would be non-aversive). During this test, both control and HI kits explored both sides (**Figure 5**). Representative traces of the path of control and HI kits are seen in Figures 5A and 5B, respectively. Control and HI kits traveled the same distance on the aversive side (t=0.7872, p=0.4359) (Figure 5C), but HI kits travelled farther on the non-aversive side (t=2.404, p=0.0204) (Figure 5D). This is partially explained by the fact HI kits spent less time on the aversive side (t=3.533, p=0.0009) (Figure 5E) and more time on the non-aversive side (t=3.533, p=0.0009) (Figure 5F) compared to control kits. In addition, when compared as a ratio, the time on the aversive: time on the non-aversive side for control kits was an average of approximately 1:1, indicating no preference for either side (Figure 5G). In contrast, the same ratio in HI kits was approximately 1:2 and was significantly different from controls (U=145.5, p=0.0016), indicating a strong preference for the non-aversive side (Figure 5G). Additionally, compared to control kits, HI kits spent a smaller percentage of their time on the aversive side immobile (U=164, p=0.0060) (Figure 5H). In contrast, HI and control kits spent the same percentage of their time on the non-aversive side immobile (t=1.217, p=0.2314) (Figure 5I). When compared as a ratio, the percent time immobile on the aversive side: the non-aversive side was approximately 1:1 in both control and HI kits, and there was no difference between the groups (U=236, p=0.2061) (Figure 5J). This indicates HI rabbits may respond to aversive textures in their environment by actively moving off of them.

**Figure 5:**
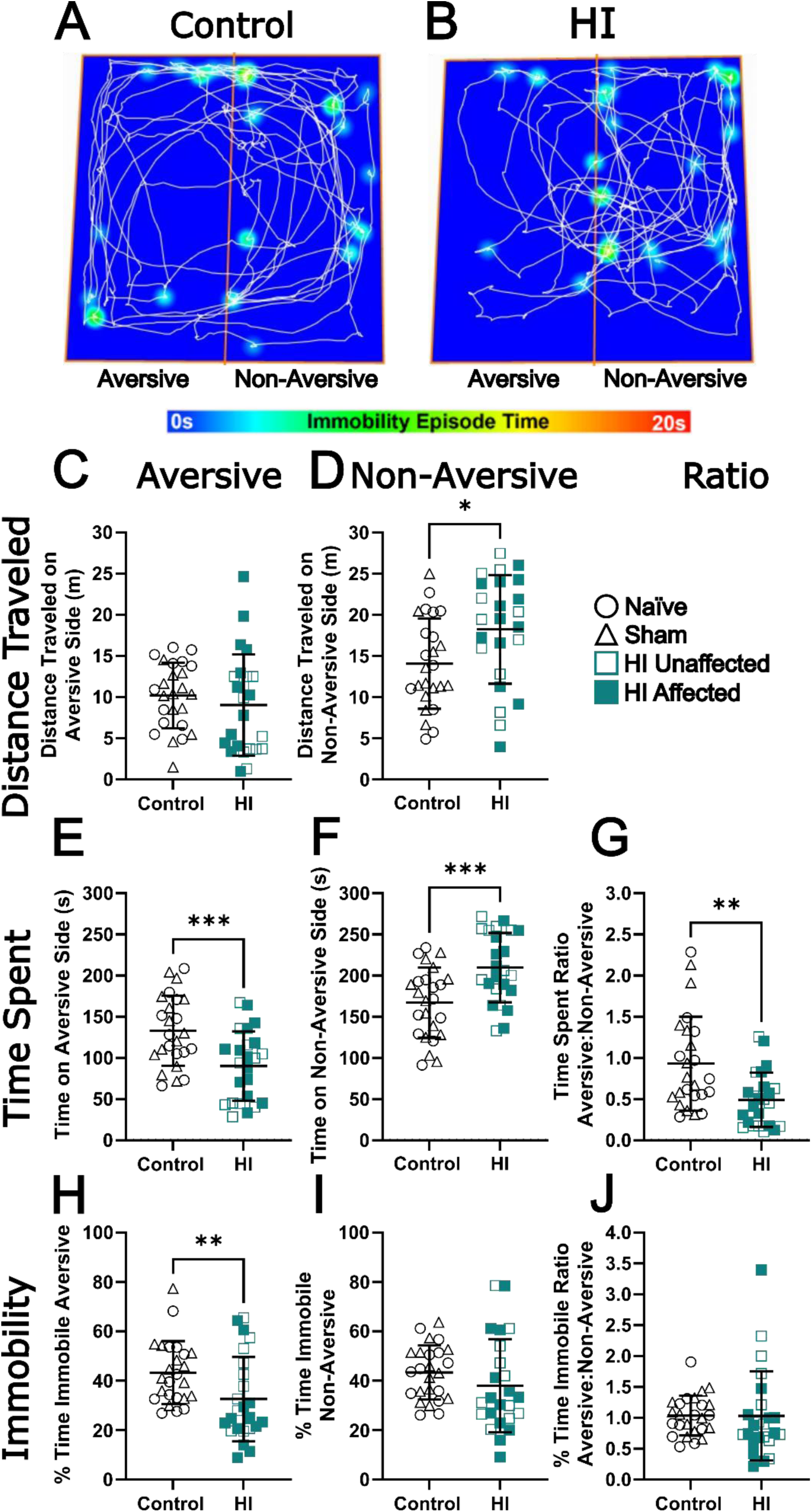
HI kits prefer the non-aversive texture during the two-texture preference test. Representative paths and immobility episodes of (**A**) control and (**B**) HI kits in the two-texture open field apparatus. (**C**) Control and HI kits traveled the same distance on the aversive side (t=0.7872, df=39.20, p=0.4359, Welch’s t test). (**D**) HI kits traveled farther than control kits on the non-aversive side (t=2.404, df=44.79, p=0.0204, Welch’s t test). (**E**) HI kits spent less time on the aversive texture as compared to control kits (t=3.533, df=46.96, p=0.0009, Welch’s t test). (**F**) HI kits spent more time on the non-aversive texture as compared to control kits (t=3.533, df=46.96, p=0.00009, Welch’s t test). (**G**) The ratio of time spent on the aversive: non-aversive side is decreased in HI kits as compared to control kits (U=145.5, p=0.0016, Mann-Whitney test). (**H**) HI kits spent a smaller percentage of their time on the aversive side immobile as compared to control kits (U=164, p=0.0060, Mann-Whitney test). (**I**) Control and HI kits spent the same percentage of their time on the non-aversive side immobile (t=1.217, df=36.49, p=0.2314, Welch’s t test). (**J**) The ratio of the percent of time spent immobile on the aversive: non-aversive side was the same in control and HI kits (U=236, p=0.2061, Mann-Whitney test). Data represented as mean +/- SD. Control N = 25 kits, HI N = 24 kits, *p<0.5, **p<0.01, ***p<0.001

Qualitatively, the majority of control rabbits explored the area freely with little hesitation. Controls appear unbothered by the aversive texture, spending, on average, close to half the time on the turf. Only a few control rabbits licked their paw or kicked their hindpaw(s) behind them when hopping, which may be inferred as nocifensive behavior. This is in stark contrast to the behavior of HI rabbits. The HI rabbits were more hesitant to stand on the turf. At the beginning of testing, many of the HI rabbits would approach the turf, place their forepaws on it, pause, and then change direction which is visualized by the number of longer periods of immobility at the line dividing aversive from non-aversive sides of the apparatus (Figure 5B). HI rabbits would often, either while on the turf or right after being on the turf, display the nocifensive behaviors mentioned above (licking or kicking). Overall, HI kits display a strong preference for the non-aversive side of the apparatus. This data demonstrates prenatal HI is associated with increased mechanical sensitivity up to P18 despite there being no differences in the von Frey test at P18.

### Prenatal HI increases context-dependent immobility in the open field at P18

To measure anxiety-like behaviors, P18 kits were allowed to explore a 1m x 1m open field. **Figures 6A** and 6B display representative paths and location/duration of immobility from control and HI rabbit kits, respectively. When in the open field, HI and control kits locomoted similarly. Both control and HI kits traveled the same distance in the center zone (t=0.5337, p=0.5963) (Figure 6C) and the surround zone (U=279, p=0.8622) (Figure 6D). HI kits spent more time in the center zone (U=180, p=0.0257) (Figure 6E) and less time in the surround zone (U=180, p=0.0257) (Figure 6F). The ratio of time spent in the center zone: the time spent in the surround zone was greater in HI kits (U=180, p=0.0257) (Figure 6G). In addition, HI and control kits spent the same percentage of time immobile in the center zone (U=213.5, p=0.1265) (Figure 6H) and surround zone (t=0.3102, p=0.7578) (Figure 6I). However, the ratio of the percentage of time immobile in the center: the percentage of time in the surrounding area was greater in HI kits as compared to control kits (t=2.349, p=0.0233) (Figure 6J). These data begin to illuminate an increase in anxiety-like behavior at P18 in HI kits that coincides with the sensory changes seen in HI kits.

**Figure 6:**
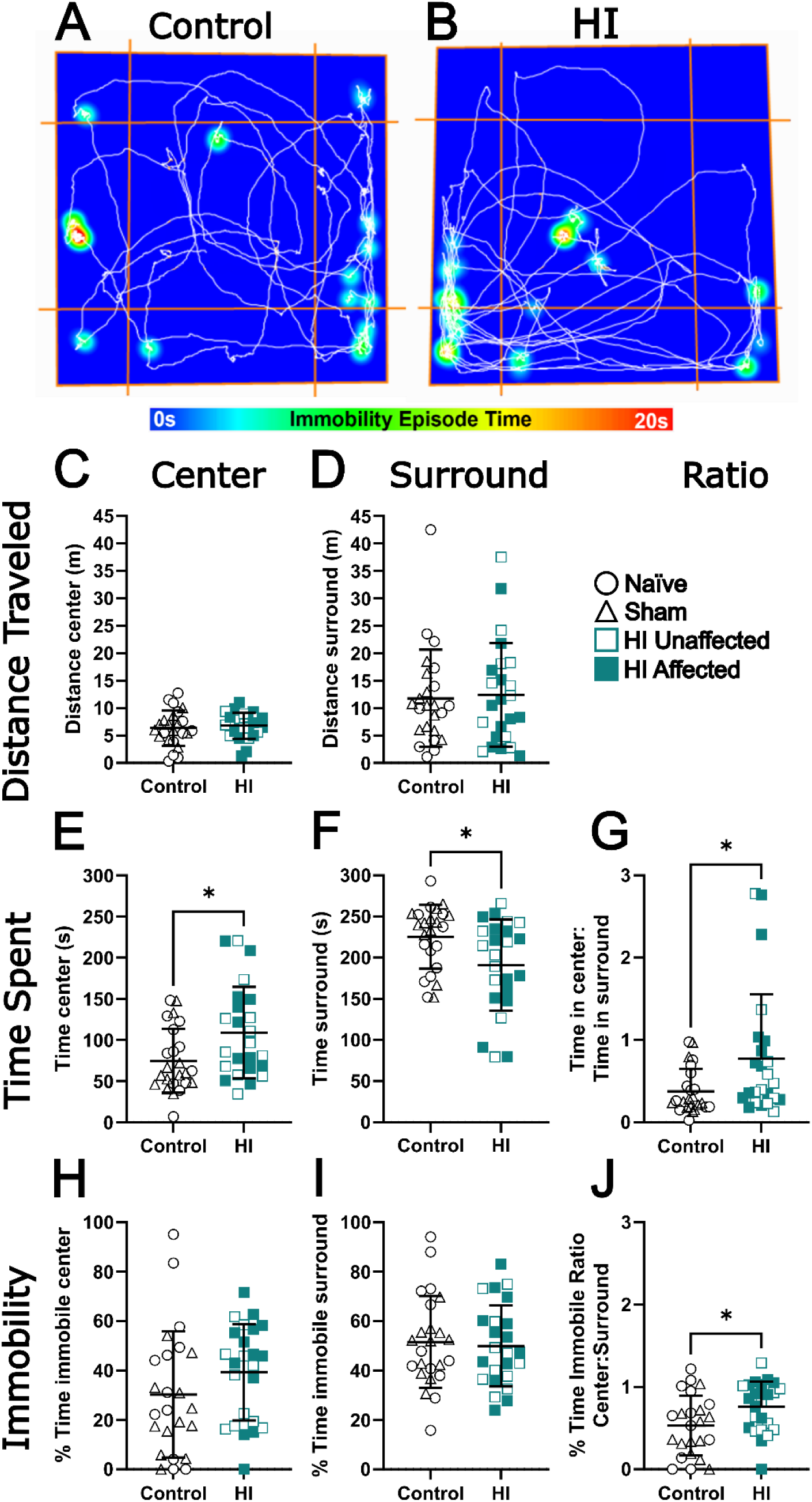
HI kits display increased anxiety-like behavior. Representative paths and immobility episodes of (**A**) control and (**B**) HI kits in the two-texture open field apparatus. (**C**) Control and HI kits traveled the same distance in the center zone (t=0.5337, df=42.15, p=0.5963, Welch’s t test). (**D**) Control and HI kits traveled the same distance in the surround zone (U=279, p=0.8622, Mann-Whitney test). (**E**) HI kits spent more time in the center zone than control kits (U=180, p=0.0257, Mann-Whitney test). (**F**) HI kits spent less time in the surround zone than control kits (U=180, p=0.0257, Mann-Whitney test). (**G**) The ratio of time spent in the center: surround zones was larger in HI kits than control kits (U=180, p=0.0257, Mann-Whitney test). (**H**) Control and HI kits spent the same percentage of their time in the center zone immobile (U=213.5, p=0.1265, Mann-Whitney test). (**I**) Control and HI kits spent the same percentage of their time in the surround zone immobile (t=0.3102, df=45.37, p=0.7578, Welch’s t test). (**J**) The ratio of time spent immobile in the center: surround zone was higher in HI kits as compared to control kits (t=2.349, df=44.67, p=0.0233, Welch’s t test). Data represented as mean +/- SD. Control N = 24 kits, HI N = 24 kits, *p<0.05

### Prenatal HI differentially alters nociceptive afferent density in the cervical and lumbar spinal cord

Nociceptive primary afferent fibers transmit pain and temperature information from the periphery. Therefore, we examined the effect of prenatal HI on nociceptive afferent distribution and density in the dorsal horn at P18. **Figure 7A-A”** shows representative images of CGRP+ and IB4+ fibers in the cervical dorsal horn of control kits. CGRP+ fibers terminate in lamina I and outer layer of lamina II, while IB4+ fibers terminate in the inner layer of lamina II in the dorsal horn of the spinal cord, with little overlap [9; 21]. Additionally, CGRP+ nociceptive afferents also terminated in the deep dorsal horn in control kits (Figure 7B-B”).

**Figure 7:**
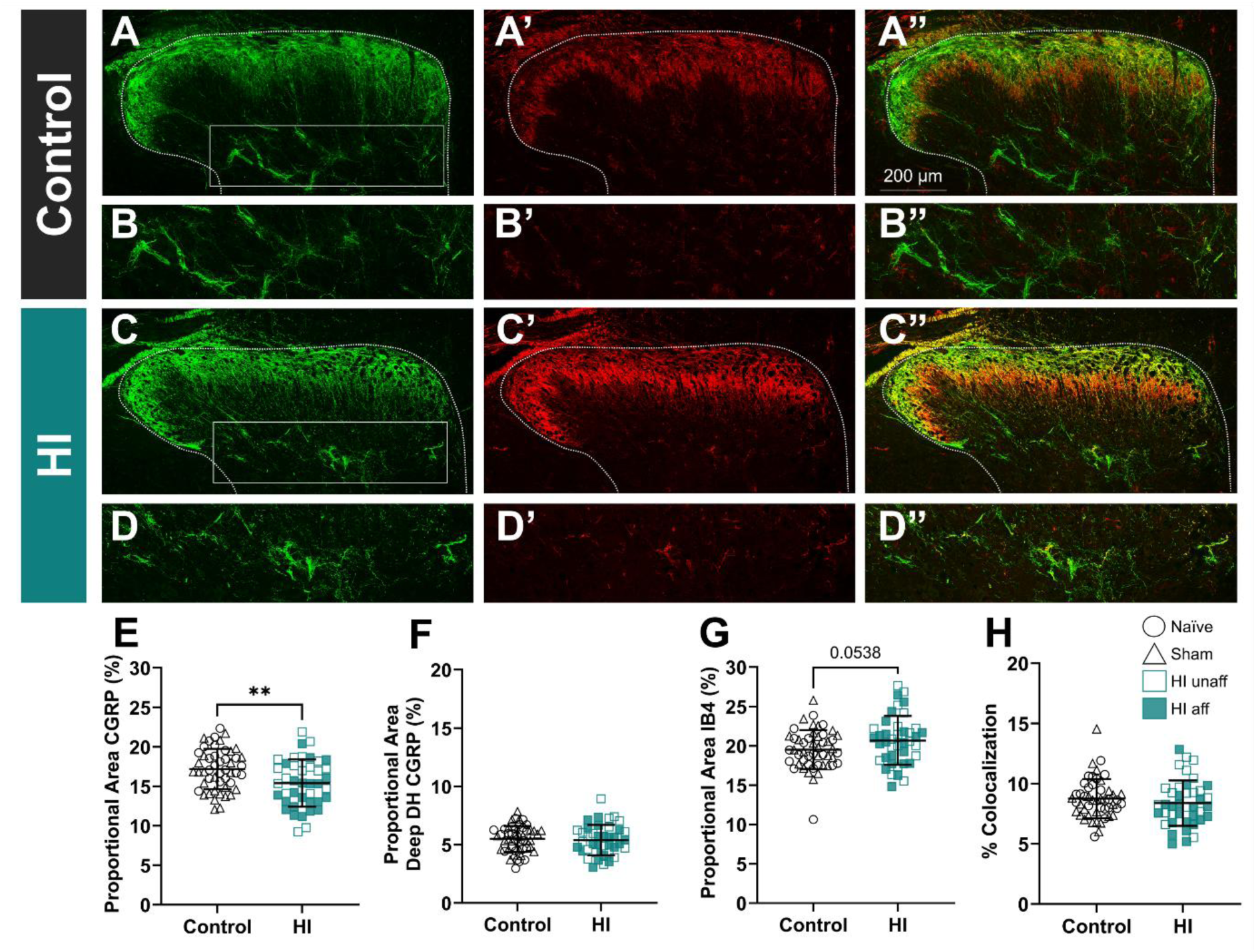
HI rabbit kits have decreased density of CGRP+ nociceptive primary afferents in the C7-8 dorsal horn at P18. Representative images of the cervical dorsal horn labeled with calcitonin gene-regulated peptide (CGRP; **A**, **B**) or labeled with isolectin B-4 (IB4; **A’**, **B’**). The proportional area of CGRP-positive tissue significantly decreased in the superficial dorsal horn in HI kits as compared to control kits (**E** t=2.991, df=79, p=0.0037, Welch’s t test). CGRP area was also measured in the deep dorsal horn (**C-C”**, **D-D”**). but there was no change between HI and control tissue (**F** t=0.6723, df=79.39, p=0.6723, Welch’s t test). There was a trending increase in the proportional area of IB4-positive tissue in HI kits as compared to control kits (**G** t=1.959, df=76.46, p=0.0538, Welch’s t test). Colocalization of CGRP-positive and IB4-positive fibers (**A”**, **B”**) was unchanged between groups (**H** U=894, p=0.2995, Mann-Whitney test). Data represented as mean ± SD; individual data points represent averages of 2-3 separate images from each left and right dorsal horn per animal. Sham and naïve kits are pooled as control kits, with circles representing naïve kits and triangles representing sham kits (n=12 naïve, n=13 sham, and n=21 HI kits). **p<0.01.

Cervical nociceptive afferents are distributed similarly in HI kits as in control kits (Figure 7C-C”, D-D”). CGRP+ nociceptive afferents terminated in lamina I, II_o,_ and III and IB4+ nociceptive afferents terminated in lamina II_i_ (Figure 7C-C”, D-D”). Surprisingly, the proportional area of superficial CGRP+ nociceptive afferents decreased after HI as compared to control kits (t=2.991, p=0.0037) (Figure 7E), but there were no group differences in CGRP+ fiber area in the deep dorsal horn (t=0.4245, p=0.6723) (Figure 7F). There was a trending increase in IB4+ fiber area in HI kits compared to controls (t=1.959, p=0.0538) (Figure 7G). In addition, peptidergic and non-peptidergic nociceptive afferent populations maintain distinct topographical distribution in both control and HI groups in the cervical dorsal horn with no changes to colocalization between groups (U=894, p=0.2995) (Figure 7H).

Lumbar nociceptive primary afferent distribution in control kits was similar to the cervical cord at P18 (**Figure 8A-B**). There was no difference between control and HI kits in the proportional area of superficial CGRP+ fibers (t=1.627, p=0.1079) (Figure 8E), but HI kits did have a significantly greater proportional area of CGRP+ fibers in the deep dorsal horn (U=537, p=0.0256) (Figure 8F), consistent with our previous finding at P5 [46], and IB4+ fibers in the superficial dorsal horn (t=2.062, p=0.043) (Figure 8G) compared to control kits. Like in the cervical dorsal horn, peptidergic and nonpeptidergic fibers maintain distinct distributions, and there are no differences in colocalization in the lumbar dorsal horn (U=786, p=0.6215) (Figure 8H). This contrasts with our previous finding of increased colocalization of CGRP+ and IB4+ fibers at P5 in HI kits [46]. The differences are likely due to the developmental trajectory. Overall, our findings demonstrate differential effects of prenatal HI on P18 nociceptive afferent area at different spinal cord levels that correspond to forelimb and hindlimb dermatomes[29; 39; 47].

**Figure 8:**
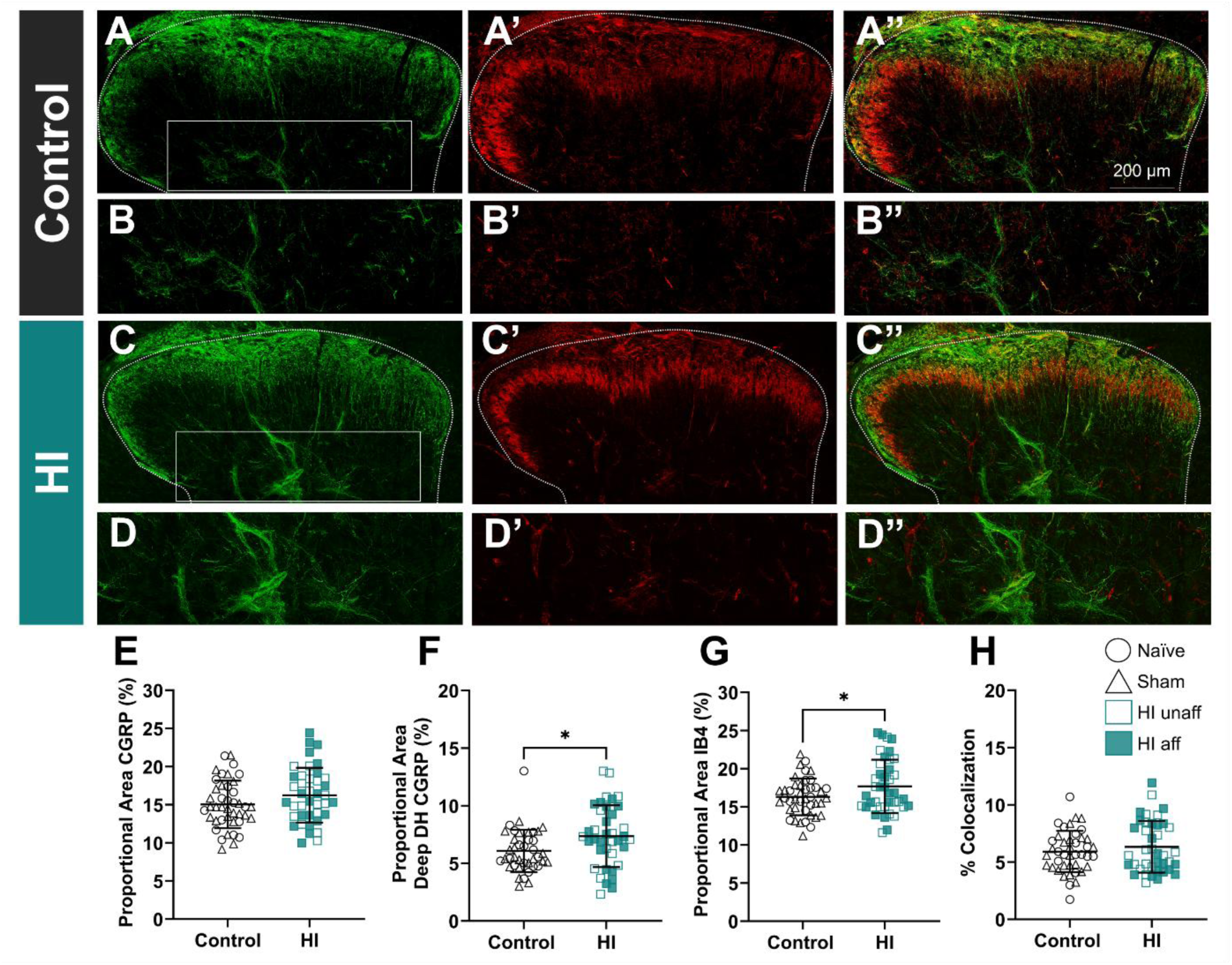
HI kits have increased density of IB4+ and deep dorsal horn CGRP+ nociceptive primary afferents in the L4-5 dorsal horn at P18. Representative images of the lumbar dorsal horn labeled with calcitonin gene-regulated peptide (CGRP; **A**, **B**) or labeled with isolectin B-4 (IB4; **A’**, **B’**). The proportional area of CGRP-positive tissue in the superficial dorsal horn remained unchanged in HI kits as compared to control kits (**E** t=1.627, df=77.26, p=0.1079, Welch’s t test). But the proportional CGRP area increased in the deep dorsal horn (**C-C”**, **D-D”**) in HI as compared to control tissue (**F** U=537, p=0.0256, Mann-Whitney test). The proportional area of IB4 positive tissue also increased in HI kits as compared to control kits (**G** t=2.062, df=68.86, p=0.043, Welch’s t test). Colocalization of CGRP-positive and IB4-positive fibers (**A”**, **B”**) was unchanged between groups (**H** U=786, p=0.6215, Mann-Whitney test). Data represented as mean ± SD; individual data points represent averages of 3 separate images from each left and right dorsal horn per animal. Sham and naïve kits are pooled as control kits, with circles representing naïve kits and triangles representing sham kits (n=9 naïve, n=12 sham, and n=21 HI kits). *p<0.05

### Principal component analysis reveals relationships between variables distinguishing control and HI kits

Principal component analysis was conducted to reduce dimensionality across the behavioral and immunofluorescence dataset. Four principal components collectively accounted for 67.8% of the total variance (**Figure 9**). The variables contributing most strongly to each PC (the absolute value of the loading coefficient ≥ 0.45) are summarized in **Table 1**.

**Figure 9:**
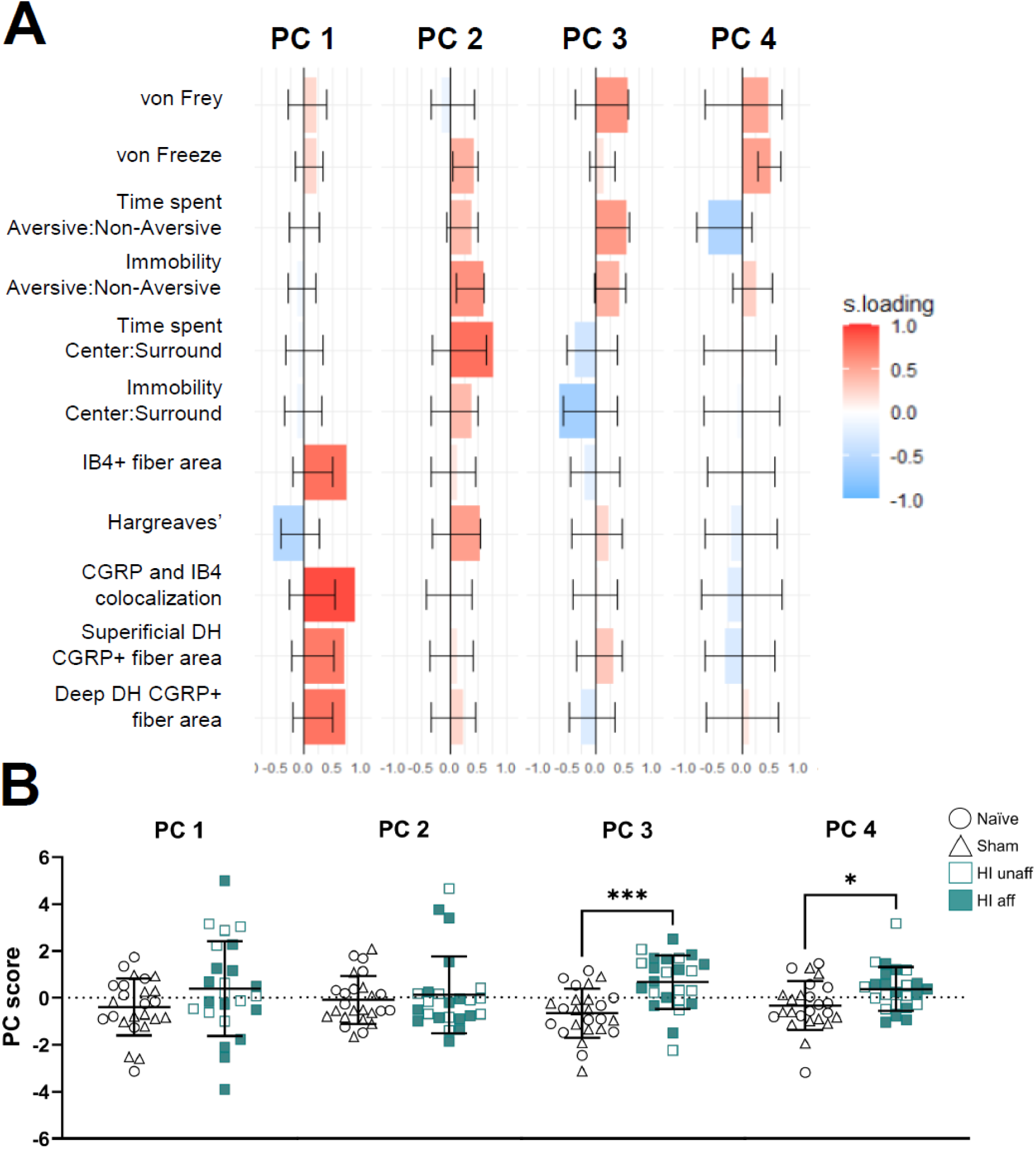
Two-texture preference, open field anxiety, and nociception variables distinguish HI from control kits. Principal component analysis revealed new uncorrelated variables called principal components (PCs). (**A**) The relative contributions of variables within PC 1-4. PC 1 has high contributions from Hargreaves’ scores and immunofluorescence variables. PC 2 has high contributions from the open field ratio of time spent in center vs surround, ratio of immobility on different textures, and Hargreaves’ paw withdrawal latencies. PC 3 has high contributions from two-texture preference ratio of time spent on each texture, open field ratio of immobility in center vs surround, and von Frey paw withdrawal thresholds. PC 4 has high contributions from von Frey paw withdrawal thresholds, von Freeze paw withdrawal latencies, and ratio of time spent on the different textures. (**B**) PC scores are plotted by experimental group. HI kits had increased PC 3 (t=4.218, df=46.25, p=0.0001, Welch’s t test) and increased PC 4 (t=2.511, df=46.78, p=0.0156, Welch’s t test) scores compared to control kits, while there was no difference between HI and control kits in PC 1 (t=1.641, df=37.41, p=0.1092, Welch’s t test) and PC2 (U=292, p=0.8819, Mann-Whitney test). Data in (B) represented as mean +/- SD. *p<0.05, **p<0.01, ***p<0.001

**Table 1:**
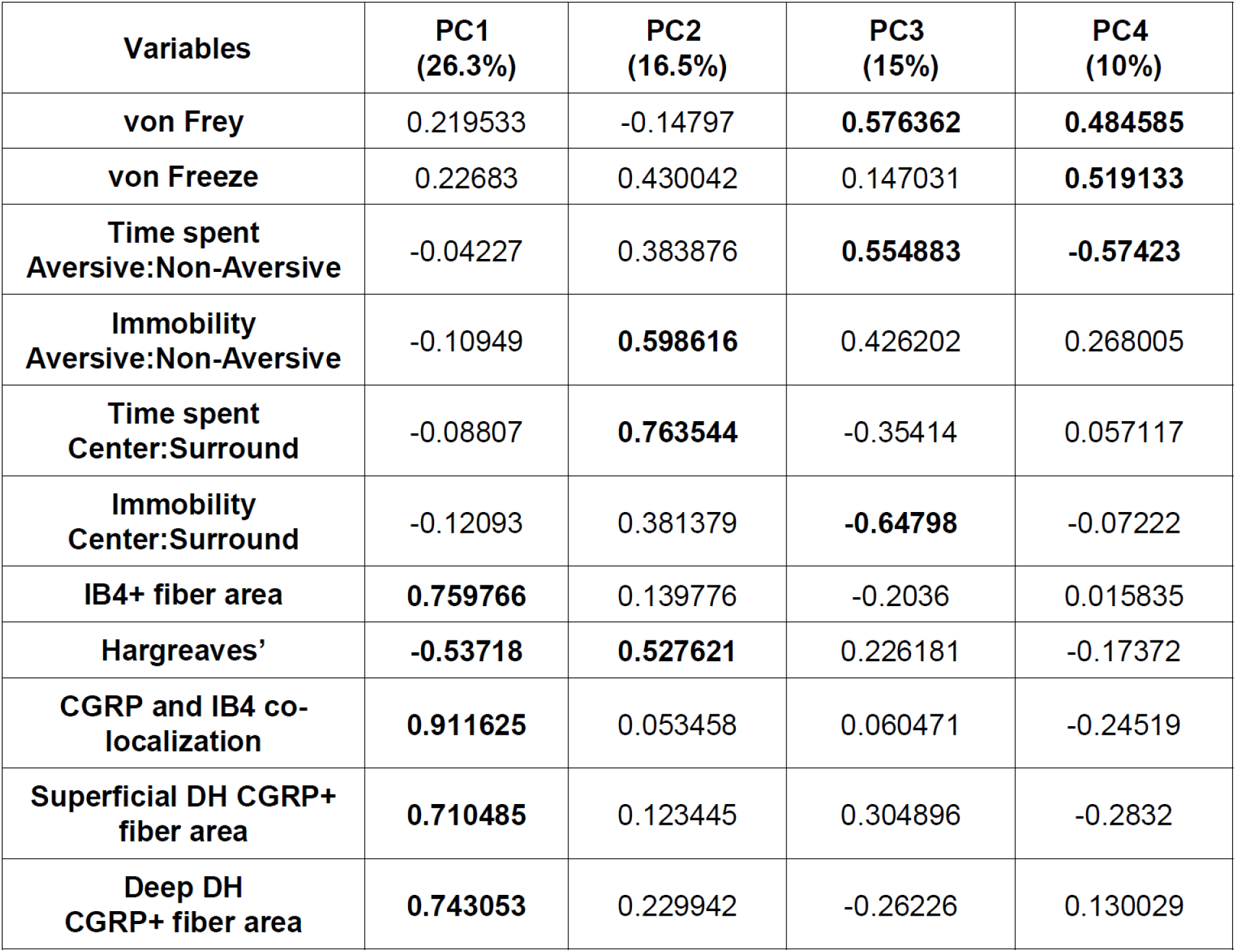
Loading values for each variable in the first four principal components.

| **Variables** | **PC1**<br>**(26.3%)** | **PC2**<br>**(16.5%)** | **PC3**<br>**(15%)** | **PC4**<br>**(10%)** |
| --- | --- | --- | --- | --- |
| **von Frey** | 0.219533 | -0.14797 | **0.576362** | **0.484585** |
| **von Freeze** | 0.22683 | 0.430042 | 0.147031 | **0.519133** |
| **Time spent**<br>**Aversive:Non-Aversive** | -0.04227 | 0.383876 | **0.554883** | **-0.57423** |
| **Immobility**<br>**Aversive:Non-Aversive** | -0.10949 | **0.598616** | 0.426202 | 0.268005 |
| **Time spent**<br>**Center:Surround** | -0.08807 | **0.763544** | -0.35414 | 0.057117 |
| **Immobility**<br>**Center:Surround** | -0.12093 | 0.381379 | **-0.64798** | -0.07222 |
| **IB4+ fiber area** | **0.759766** | 0.139776 | -0.2036 | 0.015835 |
| **Hargreaves'** | **-0.53718** | **0.527621** | 0.226181 | -0.17372 |
| **CGRP and IB4 co-**<br>**localization** | **0.911625** | 0.053458 | 0.060471 | -0.24519 |
| **Superficial DH CGRP+**<br>**fiber area** | **0.710485** | 0.123445 | 0.304896 | -0.2832 |
| **Deep DH**<br>**CGRP+ fiber area** | **0.743053** | 0.229942 | -0.26226 | 0.130029 |

PC1 was dominated by immunofluorescence variables and Hargreaves’ paw withdrawal latency, suggesting that spinal nociceptive signaling and heat sensitivity are tightly associated as a distinct biological dimension of the HI phenotype (Figure 9A). PC2 was driven primarily by immobility ratio on the different textures, anxiolytic behavior in the open field, and heat thermosensitivity. The largest contributions to PC3 were mechanosensitivity, time spent on different textures, and anxiolytic behavior in the open field. PC4 also captured mechanical sensitivity (von Frey thresholds and two-texture preference variables) loading onto this component in addition to sensitivity to cold.

When PC scores were compared between control and HI kits (Figure 9B), there were no differences between groups in PC1 (t=1.641, p=0.1092) and PC2 (U=292, p=0.8819) suggesting the individual variables loading onto each PC were not sufficient to distinguish between HI and control kits. In contrast, HI kits had significantly higher PC3 scores (t=4.218, p=0.0001) and PC4 scores (t=2.511, p=0.0156) relative to controls. The separation of HI kits from controls across PC3 and PC4 is consistent with the data presented in figures 2, 4, 5, and 6, indicating HI kits exhibit altered development of mechanical and cold nociception, increased anxiety like behavior, and actively move off aversive textures.

## Discussion

Here, we used a pre-clinical HI rabbit model of CP to longitudinally assess sensory alterations in the same rabbit kits across multiple timepoints during early postnatal development and their association with psychosocial and anatomical changes commonly associated with neuropathic pain. When tested longitudinally, we found this cohort of HI rabbits consistently displayed increased mechanical and heat-evoked thermal sensitivity and altered development of both sensory modalities. We developed a new test to measure cold-evoked thermal sensitivity and found it was decreased after prenatal HI in kits’ hindpaws. Furthermore, HI kits strongly preferred not to be on the rough texture compared to control kits and showed an increase in anxiety-like behavior aligned with psychosocial aspects of chronic pain. These behavioral changes are accompanied by an expansion of IB4+, nonpeptidergic primary afferent fibers in the lumbar dorsal horn and a trend towards a similar increase in the cervical dorsal horn in HI kits which may explain the increase in mechanical nociception in HI kits [16]. CGRP+, peptidergic primary afferent fiber distribution was more variable, depending on the region of the spinal cord, but we did recapitulate our findings at P5 that there was a greater expansion of CGRP+ fibers into the deep dorsal horn of HI kits which may contribute to the changes we see in thermosensation [16; 46]. Multivariate statistics revealed variables composed primarily of two-texture preference, open field anxiety, mechanical nociception, and cold-evoked thermal sensitivity distinguish HI from control kits.

We have expanded on our previous work showing the hypersensitivity of HI kits at P5 [46] is retained through postnatal development, and the developmental pattern of nociceptors and nociception resembles precocious development (anatomically, CGRP in the deep dorsal horn of the lumbar cord at P5 in HI resembles P18 in controls; and behaviorally, heightened mechanical nociception at younger ages in HI rabbits resembles controls at older ages). We hypothesize this is due to the brain injuries and weakened corticospinal tract underlying CP [23; 50]. With less descending control in HI rabbits, nociceptive afferents likely have less synaptic competition and dominate sensorimotor development. The range of ages that we studied in rabbits is roughly similar to children from neonatal to toddler stages based on the development of the corticospinal tract, the ability to weight support and locomote. This is consistent with literature in clinical populations showing already by ages 8 to 12, more children with CP self-report pain than the general population [44]. In a follow-up study 5 years later including many of the same participants, 74% of adolescents with CP between the ages of 13 and 17 reported pain [43]. Future studies in HI rabbits will assess behavioral and anatomical plasticity in older juvenile and adult rabbits.

Mechanical and thermal nociception was reliably altered in HI kits. These sensory modalities and the directionality of the changes may implicate changes in specific nociceptor populations or channels such as Piezo channels [63], TRPV1 channels [38] or TRPM8 channels [38]. Interestingly, Black et al. associated TRPM8-positive fibers with proprioception, suggesting a potential link between nociception and the hyperreflexia that marks spasticity [5]. Further analysis of the molecular changes that may be responsible for the hypersensitivity we observe in HI kits is warranted.

The two-texture preference test is a complex task involving cortical regions responsible for perception and execution of motor responses [2] and is a strong validation of our von Frey results at earlier ages. The difference in results between von Frey at P18 and the two-texture preference assay suggests there may be a floor effect in von Frey which kept us from detecting differences in von Frey at P18. As rabbits age, two-texture preference may be a more reliable test of mechanical sensitivity and may be a more complete assessment of pain, which can be difficult in animals. HI kits displayed nocifensive behaviors and often had immobile events at the interface of the two textures while locomoting similarly to controls indicating avoidance due to pain rather than a hesitancy to move.

The sensory changes in HI kits are accompanied by alterations in anxiety-like behavior at P18. HI rabbits spent a greater proportion of time in the center of the open field compared to controls, which would traditionally indicate reduced anxiety [33]. However, they have a greater ratio of immobility time in the center zone to immobility time in the surround zone, with some freezing behavior (no visible movement besides respiration), a behavior consistent with heightened anxiety [8; 36]. The pre-weaning age of our subjects may also account for this inconsistency, as the behavioral expression of anxiety differs between juveniles and adults [3; 12; 28; 55; 57]. In the open field, control rabbits initially displayed cautious, exploratory behavior, moving slowly and extending onto their hindpaw toes before each hop, behaviors previously characterized as “fear” or “caution” in rabbits [36]. As they acclimate, controls transitioned to faster, more confident movement, termed “boldness” [36]. HI kits showed a marked shift in this pattern. Upon placement in the center, HI kits displayed risk assessment behaviors, delaying exploration, remaining near the placement location for a prolonged period. These observations are consistent with a heightened fear response and align with the broader biopsychosocial dimensions of chronic pain.

The PCA simplifies our high-dimensional dataset into 4 principal components. We found distribution of c-fiber variables and Hargreaves’ withdrawal latency loaded onto PC1, open field time spent and two-texture preference immobility loaded onto PC2, two-texture preference time spent, open field immobility and von Frey withdrawal thresholds loaded onto PC3, and two-texture preference time spent, von Frey withdrawal thresholds, and von Freeze withdrawal latencies loaded onto PC4. These data suggest a relationship between variables which may reduce the need to collect each behavioral measure and instead rely on a reduced set of measures to distinguish between HI and control kits. For example, there may not be a need to perform von Frey if kits can be aged to P18 to perform two-texture preference, a less demanding assay.

Our findings provide high-fidelity validation of the HI rabbit model of CP and reinforce its translational alignment with the clinical condition. We demonstrate HI rabbits exhibit hypersensitivity to both mechanical and heat stimuli, with less sensitivity to cold. Cleanly separating general hypersensitivity from allodynia or hyperalgesia remains difficult; however, these sensory changes, taken together with the two-texture preference and open field data, suggest that HI rabbits may be experiencing a pain state. This interpretation aligns with the clinical literature: people with CP have been shown to exhibit mechanical hypoesthesia, thermal hypoesthesia, and mechanical hyperalgesia, and 23% of individuals with CP in the same study met criteria for allodynia [7]. Disambiguating nociception, hyperesthesia, hypoesthesia and pain within the HI rabbit model remains an important direction for future work. Our findings are also consistent with what has been reported in rodent CP models, where HI-induced allodynia and hypersensitivity on the von Frey and Hargreaves tests have been documented in an age- and sex-dependent manner and linked to cytoarchitectural changes in supraspinal regions associated with somatosensation [18; 32].

This study could be limited by the fact HI litters included mildly affected and unaffected kits, no severely affected HI kits. However, prior work did not reveal robust differences between motor-affected and motor-unaffected HI kits in mechanical and thermal sensitivity [46]. Pain is present across Gross Motor Function Classification System levels, but the painful dermatomes vary [1; 4; 35; 41; 42]. Thus, it will be important to assess nociceptive and psychosocial pain in severely affected HI rabbits in future experiments. Second, due to the size of rabbit litters and the number of tests performed on each postnatal day it was unreasonable to have one experimenter perform all the tests or keep the same experimenter across each test for one rabbit. This may have increased the variability since animals have been shown to respond differently in sensory tests depending on the experimenter [56]. We accounted for this by randomizing who performed the test across litters and timepoints. All litters and groups were treated similarly. Third, some conclusions regarding the pain experience in HI kits are limited by the reliance on segmentally mediated nociceptive withdrawal reflexes to infer pain perception. To address this, we recorded supraspinal responses during von Frey, Hargreaves, and von Freeze testing to differentiate withdrawal responses that could be reflexive from those perceived at supraspinal levels. Moreover, the two-texture preference test assessed the perceived aversiveness of a rough, astroturf like surface, and the subsequent decision to navigate off it or avoid walking on it altogether. Together these assessments provide insight into the connectivity of local spinal cord circuits to supraspinal, cortical centers involved in pain perception.

## Conclusions

In conclusion, these data emphasize the clinical relevance of the HI rabbit as a model of CP pain and point to future work following pain behaviors in these rabbits into adulthood. This work and the research that builds upon it will lead to increased awareness of pain as a comorbidity of CP, a better understanding of nociception in the context of CP, and the development of more efficacious treatments for said pain.

## Acknowledgements

Funding provided by: NIH NINDS Help End Addiction Longterm (H.E.A.L.) Initiative RFNS135580 to KAQ, MRD, and NK, R01 NS104436 (KQ), R01 NS132728 (KQ) and NIH Instrumentation grant #S10OD034206. We acknowledge generative AI (ChatGPT-OpenAI) was used to generate code, streamline/simplify code, confirm correct code use, and troubleshoot code during the use of R for linear mixed effects models. We gratefully acknowledge the input and meaningful advice on CP pain from our clinical and community partners Dr. Deborah Gaebler, Dr. Benjamin Katholi, Ashleigh Nightengale, Ross Bostick, Ann Adams, as well as technical support from Grace A. Giddings.

---

**Supplemental Table 1:**
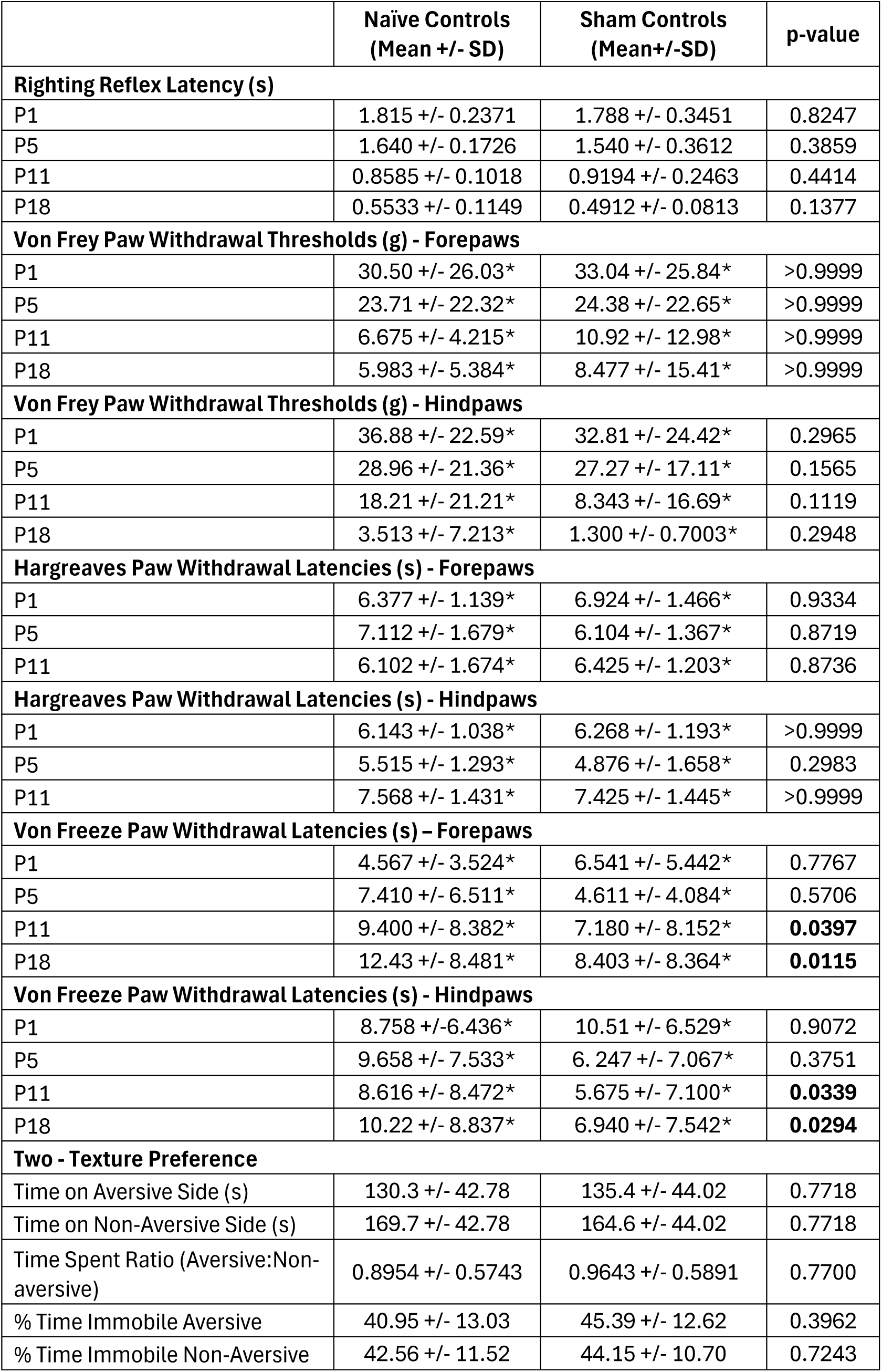
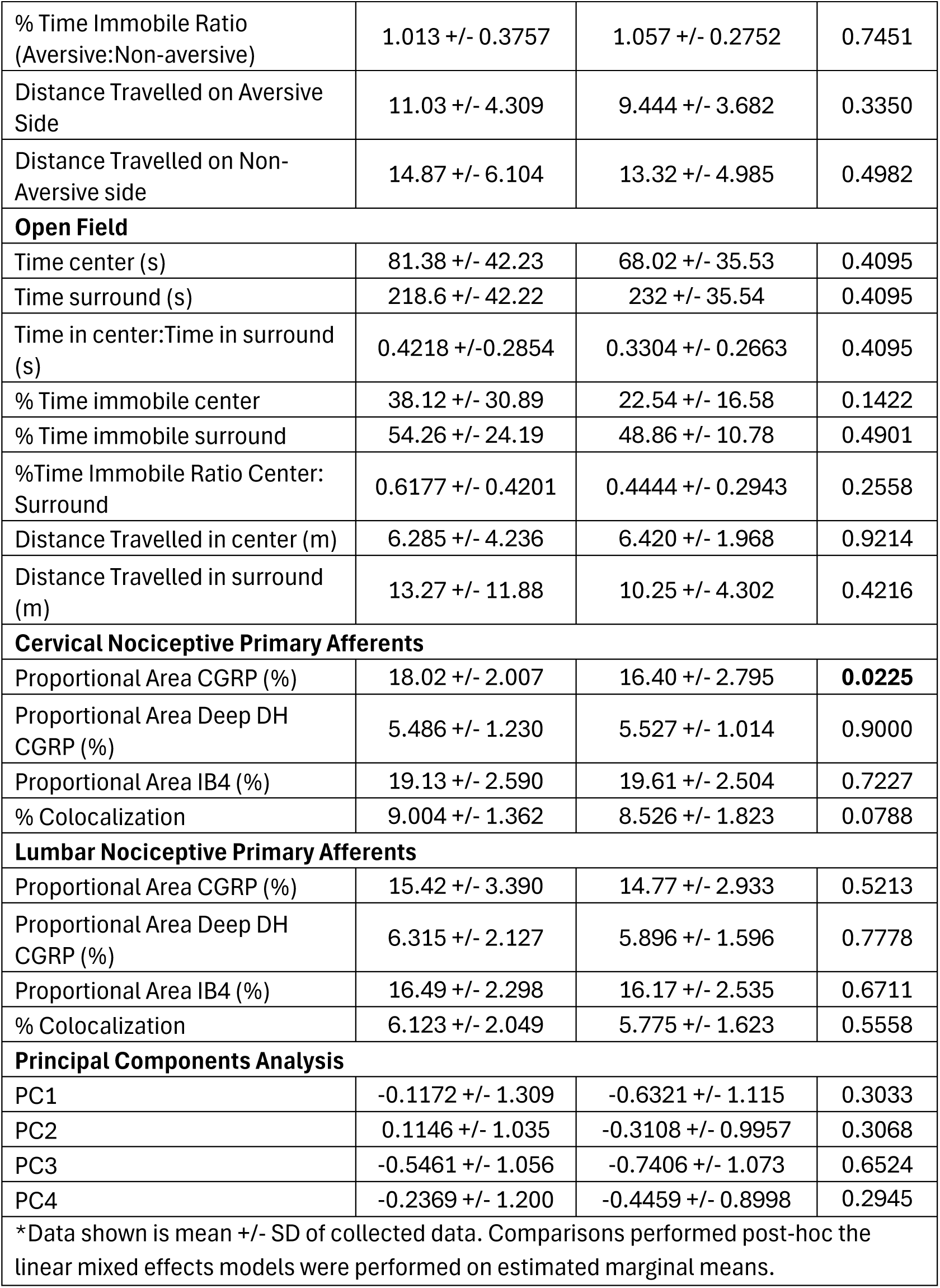
Naïve control vs sham pairwise comparisons.

**Supplemental Table 2:**
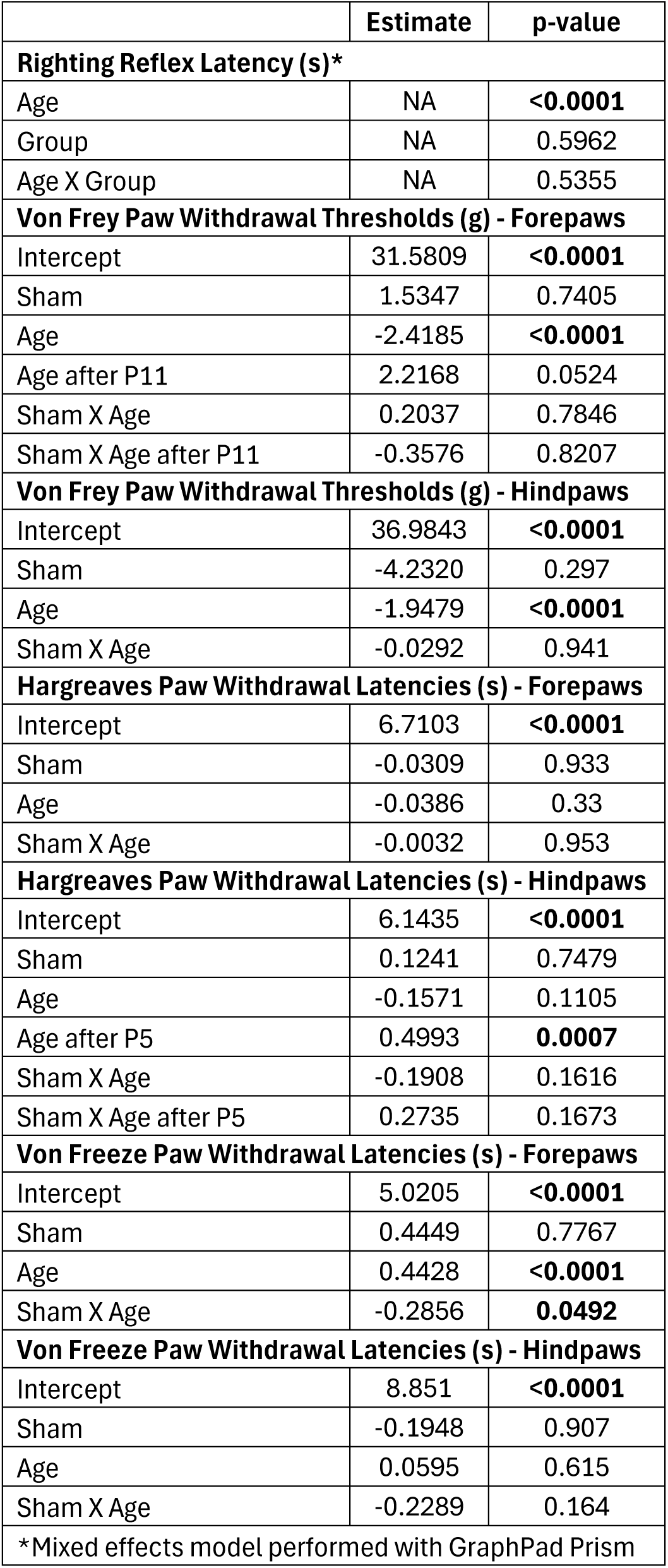
Linear mixed effects models to compare naïve controls and sham developmental trajectories in von Frey, Hargreaves, and von Freeze.

