## supplemental tables 1 and 2 for "Aberrant Pain Phenotypes Emerge Following Prenatal Hypoxic-Ischemic Injury in a Rabbit Model of Cerebral Palsy"

|  | **Naïve Controls (Mean +/- SD)** | **Sham Controls (Mean+/-SD)** | **p-value** |
| --- | --- | --- | --- |
| **Righting Reflex Latency (s)** | | | |
| P1 | 1.815 +/- 0.2371 | 1.788 +/- 0.3451 | 0.8247 |
| P5 | 1.640 +/- 0.1726 | 1.540 +/- 0.3612 | 0.3859 |
| P11 | 0.8585 +/- 0.1018 | 0.9194 +/- 0.2463 | 0.4414 |
| P18 | 0.5533 +/- 0.1149 | 0.4912 +/- 0.0813 | 0.1377 |
| **Von Frey Paw Withdrawal Thresholds (g) - Forepaws** | | | |
| P1 | 30.50 +/- 26.03* | 33.04 +/- 25.84* | >0.9999 |
| P5 | 23.71 +/- 22.32* | 24.38 +/- 22.65* | >0.9999 |
| P11 | 6.675 +/- 4.215* | 10.92 +/- 12.98* | >0.9999 |
| P18 | 5.983 +/- 5.384* | 8.477 +/- 15.41* | >0.9999 |
| **Von Frey Paw Withdrawal Thresholds (g) - Hindpaws** | | | |
| P1 | 36.88 +/- 22.59* | 32.81 +/- 24.42* | 0.2965 |
| P5 | 28.96 +/- 21.36* | 27.27 +/- 17.11* | 0.1565 |
| P11 | 18.21 +/- 21.21* | 8.343 +/- 16.69* | 0.1119 |
| P18 | 3.513 +/- 7.213* | 1.300 +/- 0.7003* | 0.2948 |
| **Hargreaves Paw Withdrawal Latencies (s) - Forepaws** | | | |
| P1 | 6.377 +/- 1.139* | 6.924 +/- 1.466* | 0.9334 |
| P5 | 7.112 +/- 1.679* | 6.104 +/- 1.367* | 0.8719 |
| P11 | 6.102 +/- 1.674* | 6.425 +/- 1.203* | 0.8736 |
| **Hargreaves Paw Withdrawal Latencies (s) - Hindpaws** | | | |
| P1 | 6.143 +/- 1.038* | 6.268 +/- 1.193* | >0.9999 |
| P5 | 5.515 +/- 1.293* | 4.876 +/- 1.658* | 0.2983 |
| P11 | 7.568 +/- 1.431* | 7.425 +/- 1.445* | >0.9999 |
| **Von Freeze Paw Withdrawal Latencies (s) – Forepaws** | | | |
| P1 | 4.567 +/- 3.524* | 6.541 +/- 5.442* | 0.7767 |
| P5 | 7.410 +/- 6.511* | 4.611 +/- 4.084* | 0.5706 |
| P11 | 9.400 +/- 8.382* | 7.180 +/- 8.152* | **0.0397** |
| P18 | 12.43 +/- 8.481* | 8.403 +/- 8.364* | **0.0115** |
| **Von Freeze Paw Withdrawal Latencies (s) - Hindpaws** | | | |
| P1 | 8.758 +/-6.436* | 10.51 +/- 6.529* | 0.9072 |
| P5 | 9.658 +/- 7.533* | 6. 247 +/- 7.067* | 0.3751 |
| P11 | 8.616 +/- 8.472* | 5.675 +/- 7.100* | **0.0339** |
| P18 | 10.22 +/- 8.837* | 6.940 +/- 7.542* | **0.0294** |
| **Two - Texture Preference** | | | |
| Time on Aversive Side (s) | 130.3 +/- 42.78 | 135.4 +/- 44.02 | 0.7718 |
| Time on Non-Aversive Side (s) | 169.7 +/- 42.78 | 164.6 +/- 44.02 | 0.7718 |
| Time Spent Ratio (Aversive:Non-aversive) | 0.8954 +/- 0.5743 | 0.9643 +/- 0.5891 | 0.7700 |
| % Time Immobile Aversive | 40.95 +/- 13.03 | 45.39 +/- 12.62 | 0.3962 |
| % Time Immobile Non-Aversive | 42.56 +/- 11.52 | 44.15 +/- 10.70 | 0.7243 |
| % Time Immobile Ratio (Aversive:Non-aversive) | 1.013 +/- 0.3757 | 1.057 +/- 0.2752 | 0.7451 |
| Distance Travelled on Aversive Side | 11.03 +/- 4.309 | 9.444 +/- 3.682 | 0.3350 |
| Distance Travelled on Non-Aversive side | 14.87 +/- 6.104 | 13.32 +/- 4.985 | 0.4982 |
| **Open Field** | | | |
| Time center (s) | 81.38 +/- 42.23 | 68.02 +/- 35.53 | 0.4095 |
| Time surround (s) | 218.6 +/- 42.22 | 232 +/- 35.54 | 0.4095 |
| Time in center:Time in surround (s) | 0.4218 +/-0.2854 | 0.3304 +/- 0.2663 | 0.4095 |
| % Time immobile center | 38.12 +/- 30.89 | 22.54 +/- 16.58 | 0.1422 |
| % Time immobile surround | 54.26 +/- 24.19 | 48.86 +/- 10.78 | 0.4901 |
| %Time Immobile Ratio Center: Surround | 0.6177 +/- 0.4201 | 0.4444 +/- 0.2943 | 0.2558 |
| Distance Travelled in center (m) | 6.285 +/- 4.236 | 6.420 +/- 1.968 | 0.9214 |
| Distance Travelled in surround (m) | 13.27 +/- 11.88 | 10.25 +/- 4.302 | 0.4216 |
| **Cervical Nociceptive Primary Afferents** | | | |
| Proportional Area CGRP (%) | 18.02 +/- 2.007 | 16.40 +/- 2.795 | **0.0225** |
| Proportional Area Deep DH CGRP (%) | 5.486 +/- 1.230 | 5.527 +/- 1.014 | 0.9000 |
| Proportional Area IB4 (%) | 19.13 +/- 2.590 | 19.61 +/- 2.504 | 0.7227 |
| % Colocalization | 9.004 +/- 1.362 | 8.526 +/- 1.823 | 0.0788 |
| **Lumbar Nociceptive Primary Afferents** | | | |
| Proportional Area CGRP (%) | 15.42 +/- 3.390 | 14.77 +/- 2.933 | 0.5213 |
| Proportional Area Deep DH CGRP (%) | 6.315 +/- 2.127 | 5.896 +/- 1.596 | 0.7778 |
| Proportional Area IB4 (%) | 16.49 +/- 2.298 | 16.17 +/- 2.535 | 0.6711 |
| % Colocalization | 6.123 +/- 2.049 | 5.775 +/- 1.623 | 0.5558 |
| **Principal Components Analysis** | | | |
| PC1 | -0.1172 +/- 1.309 | -0.6321 +/- 1.115 | 0.3033 |
| PC2 | 0.1146 +/- 1.035 | -0.3108 +/- 0.9957 | 0.3068 |
| PC3 | -0.5461 +/- 1.056 | -0.7406 +/- 1.073 | 0.6524 |
| PC4 | -0.2369 +/- 1.200 | -0.4459 +/- 0.8998 | 0.2945 |
| *Data shown is mean +/- SD of collected data. Comparisons performed post-hoc the linear mixed effects models were performed on estimated marginal means. | | | |

**Supplemental Table 1:** Naïve control vs sham pairwise comparisons

|  | **Estimate** | **p-value** |
| --- | --- | --- |
| **Righting Reflex Latency (s)*** | | |
| Age | NA | **<0.0001** |
| Group | NA | 0.5962 |
| Age X Group | NA | 0.5355 |
| **Von Frey Paw Withdrawal Thresholds (g) - Forepaws** | | |
| Intercept | 31.5809 | **<0.0001** |
| Sham | 1.5347 | 0.7405 |
| Age | -2.4185 | **<0.0001** |
| Age after P11 | 2.2168 | 0.0524 |
| Sham X Age | 0.2037 | 0.7846 |
| Sham X Age after P11 | -0.3576 | 0.8207 |
| **Von Frey Paw Withdrawal Thresholds (g) - Hindpaws** | | |
| Intercept | 36.9843 | **<0.0001** |
| Sham | -4.2320 | 0.297 |
| Age | -1.9479 | **<0.0001** |
| Sham X Age | -0.0292 | 0.941 |
| **Hargreaves Paw Withdrawal Latencies (s) - Forepaws** | | |
| Intercept | 6.7103 | **<0.0001** |
| Sham | -0.0309 | 0.933 |
| Age | -0.0386 | 0.33 |
| Sham X Age | -0.0032 | 0.953 |
| **Hargreaves Paw Withdrawal Latencies (s) - Hindpaws** | | |
| Intercept | 6.1435 | **<0.0001** |
| Sham | 0.1241 | 0.7479 |
| Age | -0.1571 | 0.1105 |
| Age after P5 | 0.4993 | **0.0007** |
| Sham X Age | -0.1908 | 0.1616 |
| Sham X Age after P5 | 0.2735 | 0.1673 |
| **Von Freeze Paw Withdrawal Latencies (s) - Forepaws** | | |
| Intercept | 5.0205 | **<0.0001** |
| Sham | 0.4449 | 0.7767 |
| Age | 0.4428 | **<0.0001** |
| Sham X Age | -0.2856 | **0.0492** |
| **Von Freeze Paw Withdrawal Latencies (s) - Hindpaws** | | |
| Intercept | 8.851 | **<0.0001** |
| Sham | -0.1948 | 0.907 |
| Age | 0.0595 | 0.615 |
| Sham X Age | -0.2289 | 0.164 |
| *Mixed effects model performed with GraphPad Prism | | |

**Supplemental Table 2:** Linear mixed effects models to compare naïve controls and sham developmental trajectories in von Frey, Hargreaves, and von Freeze.
